# The Q226H Mutation in Avian H5N1 Hemagglutinin Mediates a Path towards Structural Adaptation in Humans

**DOI:** 10.64898/2026.05.21.726965

**Authors:** Ross A. Edwards, Oluwafemi F. Adu, Egor P. Tchesnokov, Dana Kocincova, Emma Woolner, Zoe Turner, Duong T. Bui, Lara K. Mahal, Nathan Zelyas, John S. Klassen, Andrei P. Drabovich, Kalyan Das, Matthias Götte

## Abstract

The global outbreak of highly pathogenic avian influenza (HPAI) A(H5N1) among birds and the spillover to mammals increases the risk for humans. A recent case in British Columbia with a clade 2.3.4.4b H5 virus infection revealed a mixture of 226Q/H in the receptor-binding site of hemagglutinin. While significant changes in pre-existing immunity by H1 or H3 polyclonal sera are not evident, we show that the Q226H mutation enables binding to human-type a2-6 sialic acid receptors. High-resolution cryo-EM structures provide a basis for the alteration in receptor preference and show that a possible path towards human adaptation also requires a conformational change of the bound a2-6-sialylated glycan. Continued surveillance for additional mutations that could enhance this phenotype is warranted.

---

The emergence of highly pathogenic avian influenza (HPAI) A(H5N1) clade 2.3.4.4b (*1*), coupled with recent spillover events from birds to cattle and other mammals, highlights a significant pandemic potential. Sporadic H5N1 infections in humans are commonly mild, and human-to- human transmissions have not yet been reported. However, several recent cases have been associated with severe disease and raise concerns about emerging viruses with markers of adaptation (*2–6*).

In 2024, a 13-year-old adolescent in British Columbia (BC), Canada, was critically ill due to an infection with a variant of H5 clade 2.3.4.4b herein referred to as A/BC/PHL-2032/2024(H5N1) virus (A/British_Columbia/PHL-2032/2024|EPI_ISL_19548836|A/H5N1) (*2*). Sequencing revealed the E627K mutation in the polymerase basic protein 2 (PB2) (52% allele frequency) that is associated with mammalian adaptation. PB2-E627K was shown to facilitate interactions with human acidic nuclear phosphoproteins such as ANP32A or ANP32B that promote viral genome replication (*7–9*). Human adaptation is complex and involves several stages in the life cycle of influenza(*10*). Receptor specificity is an important parameter in this regard. Several recent reports provided evidence, both consistent and inconsistent with possible changes in receptor binding specificity associated with clade 2.3.4.4b viruses (*11–15*). Influenza hemagglutinin (HA) is required for receptor-mediated entry via sialic acid receptors with avian-type a2-3 specificity or human-type a2-6 specificity (*16, 17*). Avian influenza A viruses primarily recognize avian-type receptors, while adapted variants can engage human-type receptors. The A/BC/PHL-2032/2024 strain shows mixtures of HA-E190D (H3 numbering) and HA-Q226H with allele frequencies of 28% and 35%, respectively (*2*). E190 and Q226 are located in the receptor-binding site (RBS), and changes at these positions have been associated with altered receptor specificity (*17*). E190D and Q226L mutations provide prominent examples for adaptation to binding human-type sialoside linkages (*18–20*). However, the Q226H mutation has newly emerged, and its effects on the properties of HA remain unknown.

Mutations in the RBS can affect both antibody (Ab) and receptor binding (*21*). Prior infections or vaccinations with seasonal H1N1 or H3N2 viruses can provide a certain degree of protection against H5N1 infection (*22*). HA is a glycoprotein that forms trimeric complexes with variable head and conserved stem structures that allow binding of cross-reactive Abs. The RBS is in the variable head region that commonly allows binding of strain-specific, immune-dominant Abs. Binding of Abs to this region can also affect receptor binding and entry. A recent report revealed that the N169-linked glycan associated with A/Texas/37/2024(H5N1) reaches into the RBS and shows “avian-like” specificity (*23*). This “auto-glycan” adds another layer of complexity to these processes.

Here, we applied a combination of serological, biochemical and structural studies to characterize Ab, glycan and auto-glycan binding to purified A/BC/PHL-2032/2024 HA with and without the specific RBS mutations E190D and Q226H in the sialic acid-binding pocket. Polyclonal antibody (pAb) profiling with blood samples from a regional cohort of H1N1- and H3N2-infected patients reveal H5 HA stem-directed IgG1 responses with limited cross-neutralization. Cryo-EM structures of HA with H5-specific antigen-binding fragments (Fabs) of known monoclonal antibodies (mAb) confirm conserved interactions with epitopes in head and stem regions. We demonstrate that the Q226H pocket can accommodate human-type a2-6-sialylated glycans. High-resolution (∼2 Å) structures show that a conformational change of the sialic acid pyranose ring is required to avoid steric clash with the sidechain of H226. In contrast, the wild-type Q226 pocket accommodates a2- 3-sialylated glycans including the N169 auto-glycan, but not the human-type linkage. Together, these data point to a possible path towards human-type receptor binding in the context of clade 2.3.4.4b viruses.

## Cross-reactive Ab in polyclonal sera of H1 and H3 infected individuals

The recent case of critical illness with a clade 2.3.4.4b infection in British Columbia raises several questions concerning structure and function of the viral HA protein. To study possible effects on interactions with Abs or receptors, we devised two primary constructs for the expression of HA from the A/BC/PHL-2032/2024 virus: the “wild type” HA (BC24) with E190 and Q226, and the “mutant” (BC24-m) that contains the two low-frequency mutations E190D and Q226H. HA was routinely expressed in *Sf9* insect cells, while expression in mammalian Expi293F cells is specified. Guided by previous reports of pre-existing immunity to H5N1 infection (*24–26*), we initially studied the pAb response in sera or plasma samples of a small cohort from Canada, Alberta (Fig. 1). Samples of 22 individuals tested positive for H1N1 (n = 12) or H3N2 (n = 10) during the 2024– 2025 influenza season were included in this study (Table S1). Ab recognition was evaluated with the aforementioned HA proteins from BC24 and a range of contemporary H5Nx strains (Fig. S1A and B). HA sequences from A/Hong Kong/483/1997(H5N1), and recently discovered clade 2.3.4.4b viruses, including A/Astrakhan/3212/2020(H5N8) (Ast20), A/Texas/37/2024(H5N1) (TX24) and A/Washington/2148/2025(H5N5) (WA25) showed additional mutations in and outside the RBS (Fig. S1B). Purified HAs from the A/California/07/2009(H1N1) (Cal09) and A/District_Of_Columbia/27/2023(H3N2) (Col23) were used as H1 and H3 controls.

**Fig. 1.**
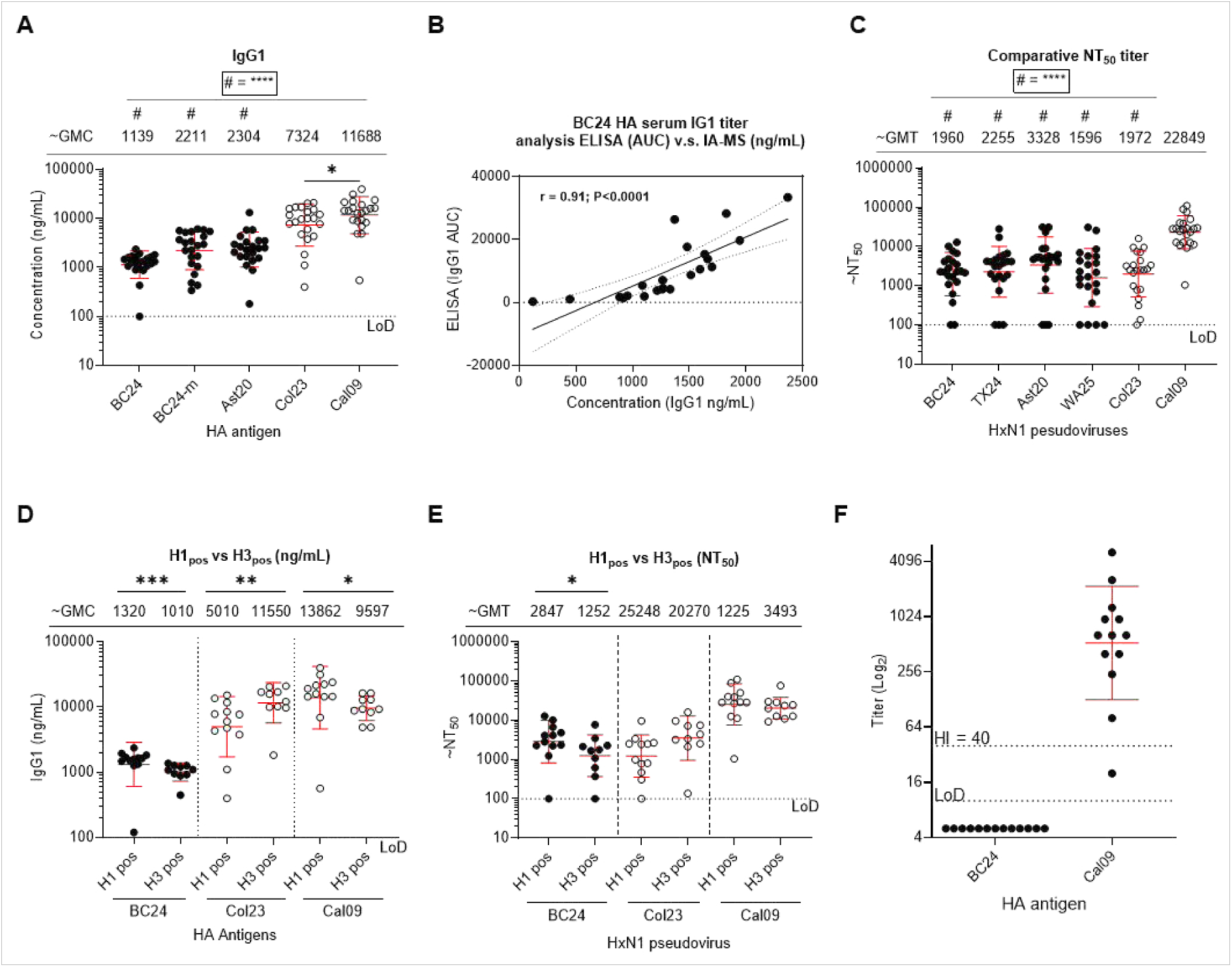
Serum Ig profile and cross-neutralization properties. (**A**) Absolute PBS-subtracted serum immunoglobulin (Ig) concentrations quantified by immunoaffinity mass spectrometry against a panel of group 1 and 2 HA recombinant antigens. Concentrations are shown as ng/mL with the geometric mean and geometric standard deviation denoted by error bars for IgG1. (**B**) Correlation plot of serum IgG1 absolute concentrations (x axis) versus area under the curve (AUC) values (y axis) derived from ELISA binding assays using the same serum samples against recombinant BC24 HA. (**C**) Serum titers represented as (NT_50_) against a panel of HxN1 lentiviruses having the same N1 from the A/BC/PHL-2032/2024 strain. Values are expressed as geometric mean titer and geometric standard deviation denoted by error bars. (**D**) Absolute IgG1 PBS-subtracted concentrations of H1N1 vs H3N2 positive individuals graphed from A. (**E**) Serum neutralization titers of H1N1 vs H3N2 positive individuals graphed from panel **D**. (**F**) HI titers, indicative of anti-head Abs recognizing BC24 HA and Cal09 HA (positive control). LoD, limit of detection (titer of 1:10) and seroprotective titer (1:40) are shown as dotted lines. All serum samples in the cohort were analyzed, while plasma samples were excluded due to anti-coagulant interference [30]. (**A-D**) groups were compared using a two-way analysis of variance (ANOVA). **** P < 0.0001, \*\*\**P* < 0.001, \*\**P* < 0.01 (E-F) H1 vs H3 positive within each group were compared using the Man-Whtiney comparison test. \*\*\**P* < 0.001, \**P* < 0.05. LoD, limit of detection (A and C) n = 2-3 biological replicates.

We employed an immunoaffinity mass spectrometry (IA-MS)-based method that enables rapid and quantitative measurement of absolute immunoglobulin (Ig) concentrations from the specimen (Fig. S2) (*27, 28*). IA-MS analysis of Abs enriched with specific antigens from serum samples allows for highly selective and absolute (ng/mL) quantification of all isotypes and subclasses of human Abs, mediated through the thoroughly designed heavy isotope-labeled quantified peptide internal standards [22]. Such analysis facilitates selection of the most informative isotypes for each antigen, and rapid screening of patient serum samples and negative controls. We first established a baseline serum Ig isotype profile using recombinant HA from the pandemic Cal09 strain (Fig. S2A-I). IgG1 dominated the response, accounting for > 94% of total anti-HA Ig, with a geometric mean concentration (GMC) of ∼11729 ng/mL, followed by IgA1 and IgM (∼526 ng/mL and ∼266 ng/mL, respectively). Other isotypes (IgG2, IgG3, IgG4, IgA2, IgD, and IgE) showed minimal reactivity above the limit of detection. Given the predominance and strongest differential response of IgG1 relative to the phosphate-buffered saline (PBS) control, subsequent analyses focused on IgG1 responses to H5N1 HA antigens (Table S2). Ab responses to the BC24, BC24-m, and reference Ast20 HAs exhibited comparable GMCs (∼1139 ng/mL, ∼2211 ng/mL, and ∼2304 ng/mL, respectively) (Fig. 1A). In contrast, responses to Col23 and Cal09 control HA proteins were substantially higher (∼7324 ng/mL and ∼11,688 ng/mL, respectively), consistent with prior exposure. These findings were validated by ELISA-based IgG1 measurements on the same sample set, which showed strong concordance with the MS data (r = 0.91), independently supporting the IgG1 response (Fig. 1B, Fig. S2J).

Serum cross-neutralization was assessed using a lentiviral pseudovirus system in a single-cycle infection format, with infection efficiency quantified via luciferase readout for viral entry (*29*). The pseudovirus panel comprised HA proteins from representative clade 2.3.4.4b H5 viruses, each paired with a common N1 neuraminidase derived from BC24. BC24-m pseudoviruses were excluded due to insufficient viral titers. GMTs were comparable across H5N1 strains (∼1960, ∼2255, ∼3328, and ∼1596 for BC24, TX24, Ast20, and WA25, respectively) (Fig. 1C). Despite these similarities, responses were heterogeneous across individuals. More than 50% exhibited NT_50_ values ≥2000, whereas a subset showed minimal cross-neutralization at or near the limit of detection. In contrast, control lentiviruses bearing Cal09 HA exhibited markedly higher titers (GMT ∼22,849), whereas those bearing Col23 HA showed titers (GMT ∼1,972) similar to the H5N1 strains. This difference may reflect imprinting, whereby early-life exposure to group 1 influenza viruses biases neutralizing responses toward H1-like epitopes in this cohort (*26*). Neutralization titers positively correlated with Ab binding to HA from BC24 (r = 0.61, Fig. S2K) suggesting that higher Ab levels contribute to increased neutralization. Stratification by infection status revealed that H1N1-infected individuals displayed higher HA-binding and cross- neutralizing responses to BC24 than H3N2-infected individuals (Fig. 1D, E), consistent with the closer phylogenetic relationship between H1 and H5 (group 1) compared to H3 (group 2) HAs (*30*).

We performed hemagglutination inhibition (HI) assays in an attempt to identify the presence of immunodominant, head-binding Ab (Fig. 1F). We determined HI titers in all serum samples of our cohort (Fig. S3). None of the tested samples exhibited significant HI activity against BC24 HA (GMT < 10). This is consistent with recent findings showing that levels of anti-head H5N1 2.3.4.4b Abs in unexposed populations are below the limit of detection in this assay (*31*). In contrast, Cal09 HA, used as a positive control, elicited robust responses (GMT 530 HI titer), exceeding the established seroprotective threshold (40 HI titer) (*32*). These findings indicate that the cross- reactive Abs found in this cohort are not primarily directed against the BC24 HA head domain.

## Binding of H5 mAbs to BC24 HA

For structural studies, we used mAbs that were previously characterized in complexes with other H5N1 HAs (Fig. 2). To assess their binding properties to both BC24 and BC24-m HAs, we expressed and tested five H5 HA head-specific mAbs (100F4, 65C6, H5.3, AvFluIgG03 and FLD21.140) (*33–35*) along with MEDI8852, a broadly neutralizing anti-stem Ab (*36*). As assayed by ELISA, 100F4, 65C6, and FLD21.140 and MEDI8852 recognized the HAs (Fig. S4). None of the Abs showed reduced binding to BC24-m HA, indicating that the RBS mutations E190D/Q226H have minimal impact on these interactions. Notably, the binding of FLD21.140 to BC24-m was retained, even though residue E190 is in contact with this Ab (*33–35*).

**Fig. 2.**
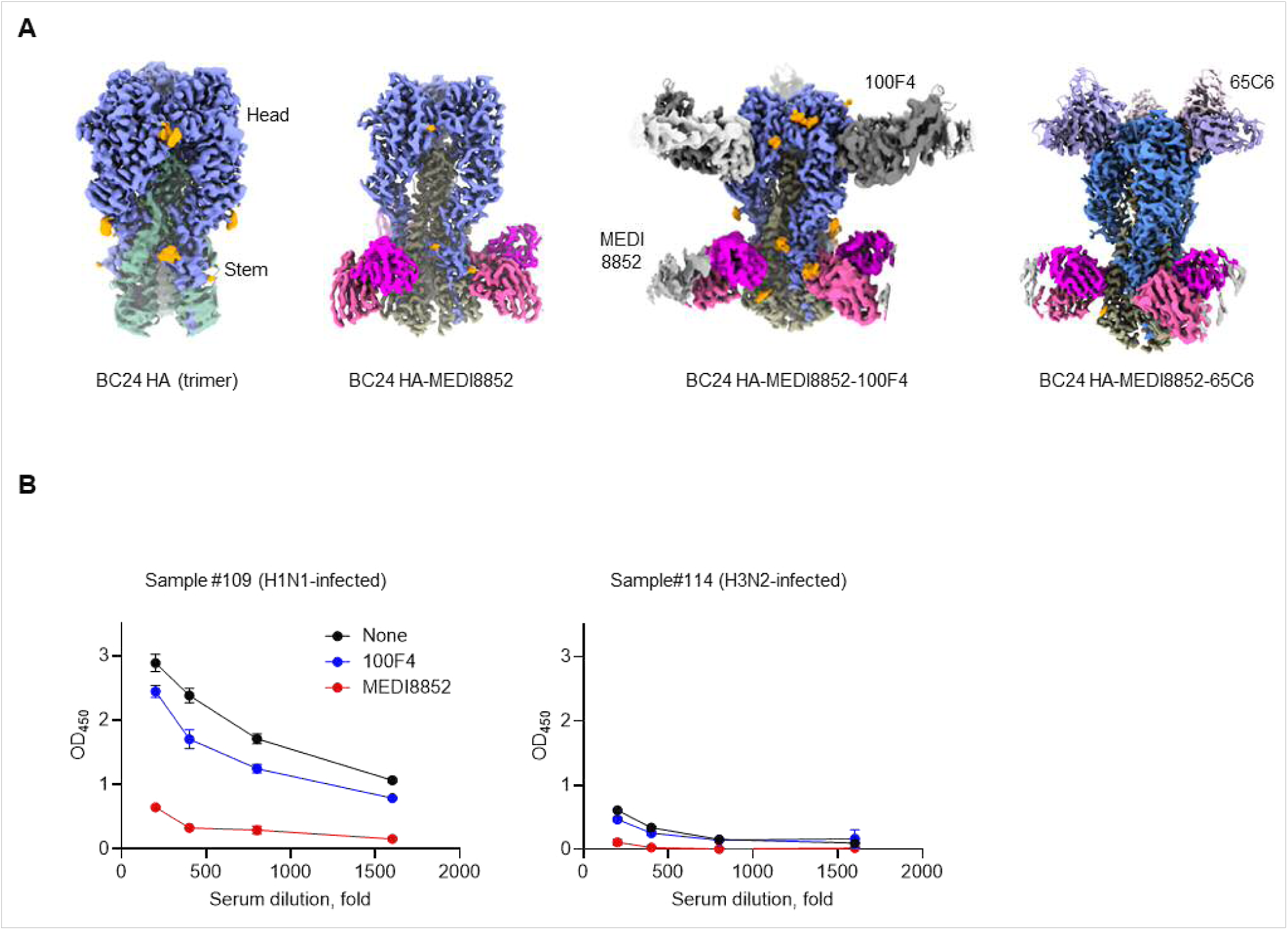
Ab binding to BC24 HA. (**A**) Cryo-EM structures of apo HA, HA in complex with mAb MEDI8852, HA in complex with MEDI8852 and 100F4, and HA in complex with MEDI8852 and 65C6 in a sequence from left to right. MEDI8852 binds to the stem, while 100F4 and 65C6 bind to the head. These are the structures I, II, IV, and VI from the eight structures reported in this study (Fig. S5). (**B**) In a competitive ELISA, BC24 HA was pre-incubated separately with no Ab, 100F4 or MEDI8852 Fabs before adding sera dilutions from H1N1- and H3N2-infected individuals, respectively. For he high signal is indicative for pAb binding and the low signal indicates either competition with mAbs or inefficient binding.

Fab fragments of the broadly neutralizing stem-binding mAb MEDI8852 were complexed with BC24 HA in combination with either of the two head-binding mAbs (*33–36*), 100F4 and 65C6. We determined eight cryo-EM structures (**I** – **VII**) of BC24 and BC24-m HAs and their complexes with Abs and glycans (Fig. S5). Structures (**I**) BC24, (**II**) BC24 with MEDI8852 mAb, (**IV**) BC24- m HA with MEDI 8852 and 100F4 (**VI**) BC24-m with MEDI8852 and 65C6, reveal binding characteristics of the three mAbs to HA (Fig. 2A). The structures are similar to those reported earlier with different H5N1 HAs (*33–36*) (Fig. S6). As expected, the mAb MEDI8852 binds the stem region, which is highly conserved in H5 HAs. The mAbs 100F4 and 65C6 bind to two distinct regions of the head. Binding of 100F4 with BC24 is somewhat altered compared to the structure of 100F4-bound A/Anhui/1/2005(H5N1) HA (*37*). Sequence differences at the interface positions 81, 122, and 173 may account for perturbed binding. Binding of 65C6 is more conserved when compared to its binding to TX24 HA (PDB 9EKF) (*23*).

Based on this data, we used the anti-head (100F4) and the anti-stem (MEDI8852) mAbs to devise a competitive ELISA to distinguish between head- and stem-binding Abs in representative H1N1- and H3N2-positive polyclonal sera (Fig. 2B, C) Pre-incubation with stem-binding MEDI8852 markedly reduced the ELISA-signal to 16% and 10% for H1N1- and H3N2-infections, respectively. In contrast, competition with the head-binding 100F4 mAbs resulted in a more subtle signal reduction, to 74% and 92% (for H1N1 and H3N2-infections, respectively). These results are complementary to the HI experiments and demonstrate that pAb responses to BC24 HA are predominantly mediated through recognition of conserved stem epitopes.

## Structural basis for binding of HA to glycan receptors

We used the Catch-and-Release native mass spectrometry (CaR-nMS) screening assay to examine glycan binding of BC24 HA and its mutant BC24-m (*38–40*). In this assay, we included forty sialylated glycans with α2-3, α2-6 and α2-8 linkages, including six complex *N*-glycans (NG1 – NG6) and thirty-four other oligosaccharides (Fig. 4A, Fig. S7). The glycans were tested against BC24 and BC24-m HAs; Cal09 HA was used as a reference, which showed a clear preference for human-type α2-6-sialylated glycans. BC24 HA showed affinity towards avian-type α2-3- sialylated glycans, and almost no binding of α2-6-sialylated glycans, in keeping with its avian origin. Most importantly, BC24-m enabled binding of α2-6-sialylated glycans (Fig. 3A, Fig. S7). The affinities of several glycans to the HAs were estimated using the Concentration Independent (COIN-nMS) method (*41*). As a negative control, α2-8-sialylated glycans showed no binding to any of the three HAs (Fig. S7).

**Fig. 3.**
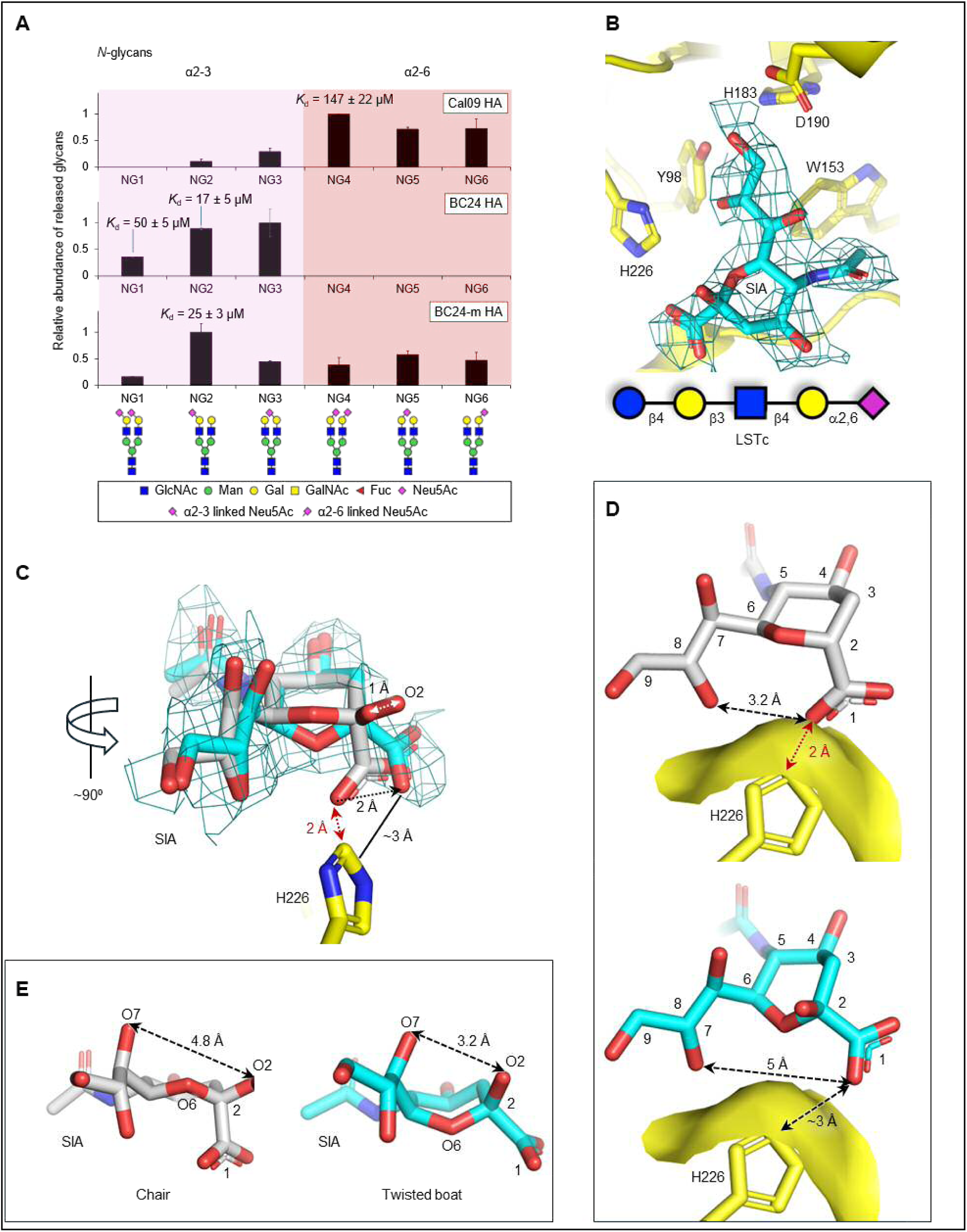
Binding of α2-6-linked sialoglycans to BC24 and BC24-m HAs. (**A**) Results of Catch-and-Release native mass spectrometry (CaR-nMS) screening of six *N*- glycans against Cal09 (top), BC24 (middle), and BC24-m (bottom) HAs. Also shown are *K*_d_ values obtained from Concentration Independent (COIN)-nMS measurements for a subset of *N*- glycan-HA interactions. The left three columns are α2-3-, and right three are analogous α2-6- sialylated *N*-glycans. As expected, the control Cal09 HA selects α2-6- and BC24 HA with E190/Q226 selects α2-3-sialylated glycans. The BC24-m HA with D190/H226 acquires some affinity to α2-6-sialylated glycans. (**B**) Binding of the sialic acid of LSTc to the H226-pocket of BC24-m HA as defined by the 1.96 Å resolution experimental density map, displayed at 4σ. (**C**) Superposition of the H5 L226-pocket-bound sialic acid of LSTc (gray, PDB 9DIO) reveals steric hinderance of 1-carboxylic group with H226, if the pyranose ring is in canonical chair conformation. In the BC24-m bound LSTc structure, the ring binds to the H226-pocket by switching its conformation to overcome the steric conflict. (**D**) The conformational switching of the pyranose ring from gray (top) to cyan (bottom), is essential for binding of sialic acid to the pocket; the C-atoms of sialic acid are numbered. (**E**) The linking O2 position is significantly shifted due to the change in the ring conformation suggesting that the position of the linked sugar rings of an α2-6-sialylated glycan should be impacted by the sialic acid conformational switching.

**Fig. 4.**
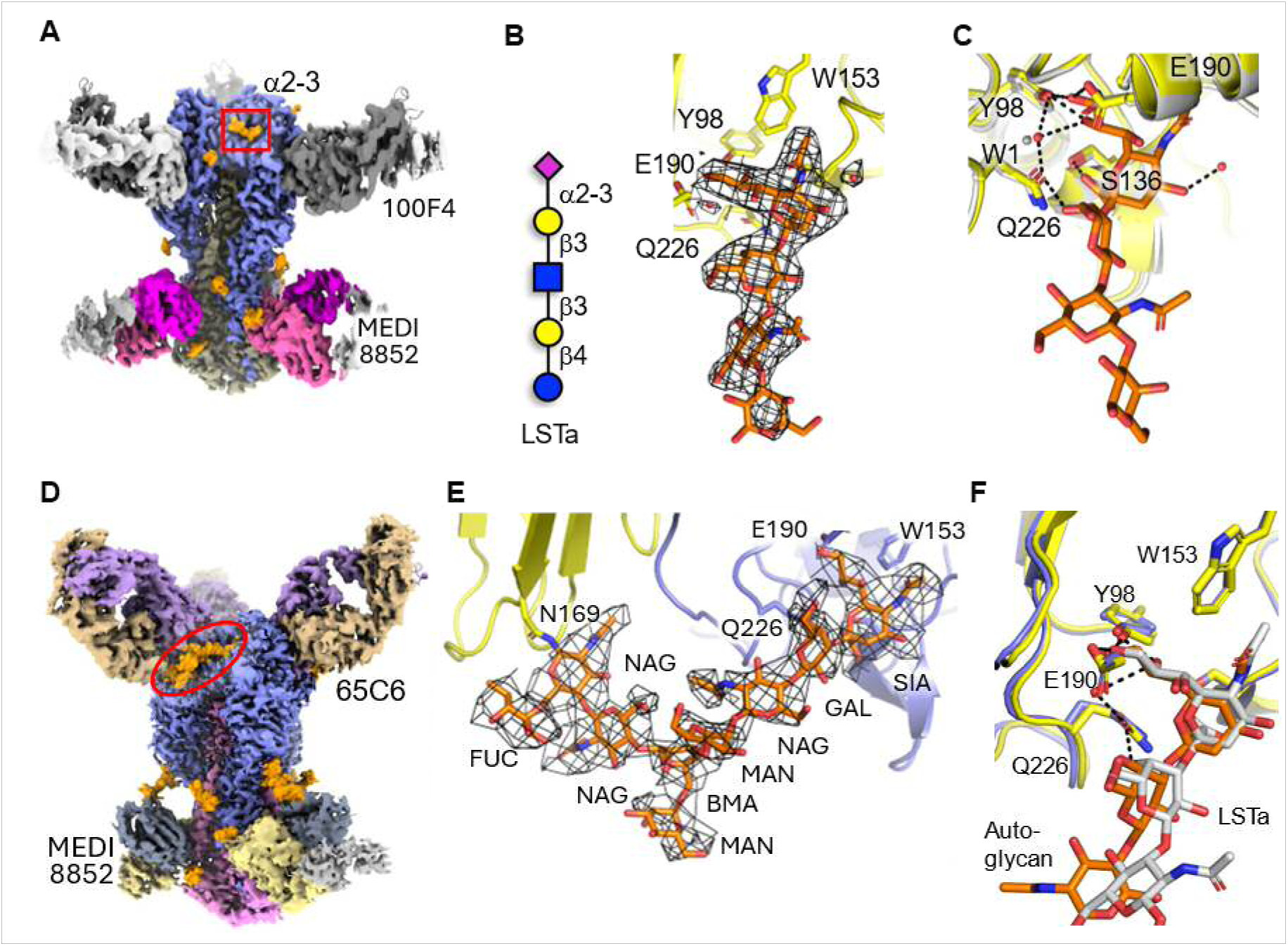
Binding of LSTa and N169 auto-glycan to BC24 HA. (**A**) The structure of BC24 HA in complex with Abs MEDI8852 and 65C6 in complex with LSTa (Fig. S5, structure **III**); LSTa (orange) binding site is indicated by a red box. (**B**) A fragment of 2.25 Å resolution cryo-EM density map displayed at 4.5σ defines all four sugar rings of LSTa. (**C**) Superposition of structure **IV** (gray) BC24 HA with an empty pocket shows no significant change in the specific pocket residues including a water molecule (W1) upon LSTa binding. (**D**) Structure **VII** of Expi293F cell- expressed BC24 HA in complex with the Abs MEDI8852 and 65C6 shows N169-glycosylated auto-glycan occupying the receptor-binding pocket (red circle). (**E**) A zoomed view of the region reveals that N169-glycosylated branched oligosaccharide with an α2-3- sialylated end occupies the pocket. The 2.85 Å resolution cryo-EM density displayed at 3σ defines the track and interactions of the auto-glycan. (**F**) The superposition of structures **III** and **VII** shows that the glycan compositions can redefine the positioning of linker sugar rings even though the interaction of the sialic acid tip is conserved.

To visualize the binding of avian-like α2-3-sialylated and human-like α2-6-sialylated glycans, we determined cryo-EM structures with model receptors LS-Tetrasaccharide a (LSTa; an α2-3- sialylated tetrasaccharide, L26 in Fig. S5) and LSTc (an α2-6-sialylated tetrasaccharide (*19*), L14), respectively. We incubated the substrates with BC24-m bound to mAbs MEDI8852 and 100F4 and determined the respective structures **IV** and **V** (Fig. S5). The structures of the mAb complexes improved the quality and resolution of the cryo-EM density maps without interfering with the glycan binding. Structure **V** of the BC24-m HA in complex with LSTc was obtained at 1.96 Å resolution (Fig. 3; Table S3). The cryo-EM density revealed binding of the sialic acid part of human-type LSTc to the receptor-binding pocket of BC24-m HA (Fig. 3B).

We observed a mutation-induced structural change in the pocket and conformational adaptation of the bound sialic acid. The Q226H mutation altered the binding pocket shape by introducing the larger aromatic imidazole side chain, which has a profound effect on substrate binding. Previously, the Q226L mutation introduced in TX24 HA was shown to cause a switch in receptor specificity from avian to human (*19*). In contrast, the Q226H mutation would not permit the binding of sialic acid in its commonly observed lowest energy “^2^C_5_ chair” conformation. This is due to steric hindrance between the H226 side chain and the C1-carboxyl group, which prevents binding of the sialic acid in a chair conformation (Fig. 3C, D). Instead, we observed a conformational change of the pyranose ring to a “twisted-boat” that positions the C1-carboxyl group away from the H226 side-chain, permitting binding of the sialic acid to the mutant pocket. This conformational change has an energetic penalty of ∼5 – 10 kcal/mol, suggesting that an α2-6-sialylated glycan would bind to the H226 pocket with significantly lower affinity than an L226 pocket. The remaining residues in the H226 pocket are minimally impacted by the Q226H mutation or the conformational switching of the sialic acid, indicating that the pocket is not yet highly adapted for improved binding of α2-6-sialylated glycans in a non-^2^C_5_ chair conformation. The conformational switching of the pyranose ring also repositioned the linking O2 atom (Fig. 3E), when compared to the TX24 HA-bound LSTc (*19*), and the O2 repositioning would impact the positioning of the remaining parts of an oligosaccharide. In our structure, the parts of LSTc beyond the sialic acid were not traceable in the density map.

We were unable to observe binding of LSTa to the RBS of the mutated HA, although Structure **IV** of BC24-m HA was obtained at 1.95 Å resolution. These findings are consistent with a diminished binding affinity of the mutated HA to α2-3-sialylated glycans. In contrast, Structure **III** of the “wild type” BC24 showed unambiguous binding to LSTa, as defined by the experimental density (Fig. 4A-C). The second pocket mutation E190D is known to facilitate human adaptation (*42*). However, we could not structurally confirm if this mutation plays a significant role in combination with Q226H mutation.

## Auto-glycosylation

Recent cryo-EM studies of TX24 HA, expressed in Expi293F cells, visualized an avian-type α2- 3-sialylated auto-glycan from N169 that occupies the RBS of the neighboring protomer (*23*). Although N169 is also glycosylated in other H5 clade 2.3.4.4b HAs, the auto-glycan was not observed occupying RBS in several other studies (*14, 19*). To address this issue, we expressed the BC24 HA in *Sf9* insect cells and Expi293F mammalian cells in parallel (Fig. S8). We observed that an α2-3-sialylated auto-glycan spans from N169 and occupies the RBS in Expi293F cell- expressed BC24 HA only (Fig. 4D-F). In the structure of Expi293F cell-expressed apo BC24 HA trimer (Fig. S5, **VII**), the N169 auto-glycan binds in a mode similar to that seen in the structure of 65C6- and MEDI8852-bound BC24 HA (Fig. S5, **VIII**). Hence, binding of mAb 65C6 adjacent to the N169 auto-glycan does not interfere with the glycan positioning. The N169 auto-glycan of HA expressed in *Sf9* cells is chemically distinct and does not occupy the RBS (Fig. S5, **I** and **II)**. Collectively, the structural data show that the pocket of BC24-m HA favors human-type α2-6- sialylated glycans and disfavors binding of avian-type α2-3-sialylated glycans, and this would include the auto-glycan from N169.

## Discussion

The global spread of HPAI H5N1 clade 2.3.4.4b into mammals alongside sporadic human infections necessitates a detailed understanding of emerging patterns of adaptation (*1*). Mutations in influenza HA are of particular concern as structural changes in this glycoprotein can alter receptor-binding specificity and/or affect Ab binding (*16, 19*). Two HA mutations, E190D and Q226H, are in the sialic acid-binding pocket. While E190D was shown to improve human-type receptor use (*20, 43*), Q226H is a newly described mutation at a position that is also known to affect adaptation (*2, 14, 15*). The Q226L mutation was recently shown to switch binding specificity to human α2-6 sialosides when introduced into clade 2.3.4.4b viruses (*19, 44, 45*). These observations highlight the need for mechanistic investigations that focus on changes in structure and function of the BC24 glycoprotein with and without pocket mutations.

We showed that the pAb response associated with seasonal H1 or H3 infections, in particular the IgG1 subclass, is likely a component of the pre-existing immunity against H5 viruses. The relevant Abs bind predominantly to the conserved stem region, which is exemplified by high-resolution cryo-EM structures of BC24 HA and BC24-m HA with MEDI8852. Most importantly, BC24-m HA has a profound impact on glycan binding. The Q226H mutation changes the pocket in a distinct manner that was previously not seen with Q226L. Consequently, the ring conformation of the sialic acid receptor must be changed to avoid a steric clash with the side chain of H226 (Fig. 5A, B). The α2-6 linkage with three rotatable bonds introduces the required flexibility that permits sialic acid binding to the H226 pocket (Fig. 5C). However, the bound LSTc is disordered and has no noticeable interaction beyond its sialic acid moiety. While the collective data do not support a bona fide receptor switch associated with the Q226H mutation, the structures demonstrate plasticity of the RBS region and its substrate that could pave the way for new adaptation mechanisms. Additional mutations at variable neighboring locations could optimize the RBS in the mutational context of Q226H (Fig. 5E, F) (*46*). Together these findings warrant detailed surveillance efforts with attention to changes that could further enhance this phenotype.

**Fig. 5.**
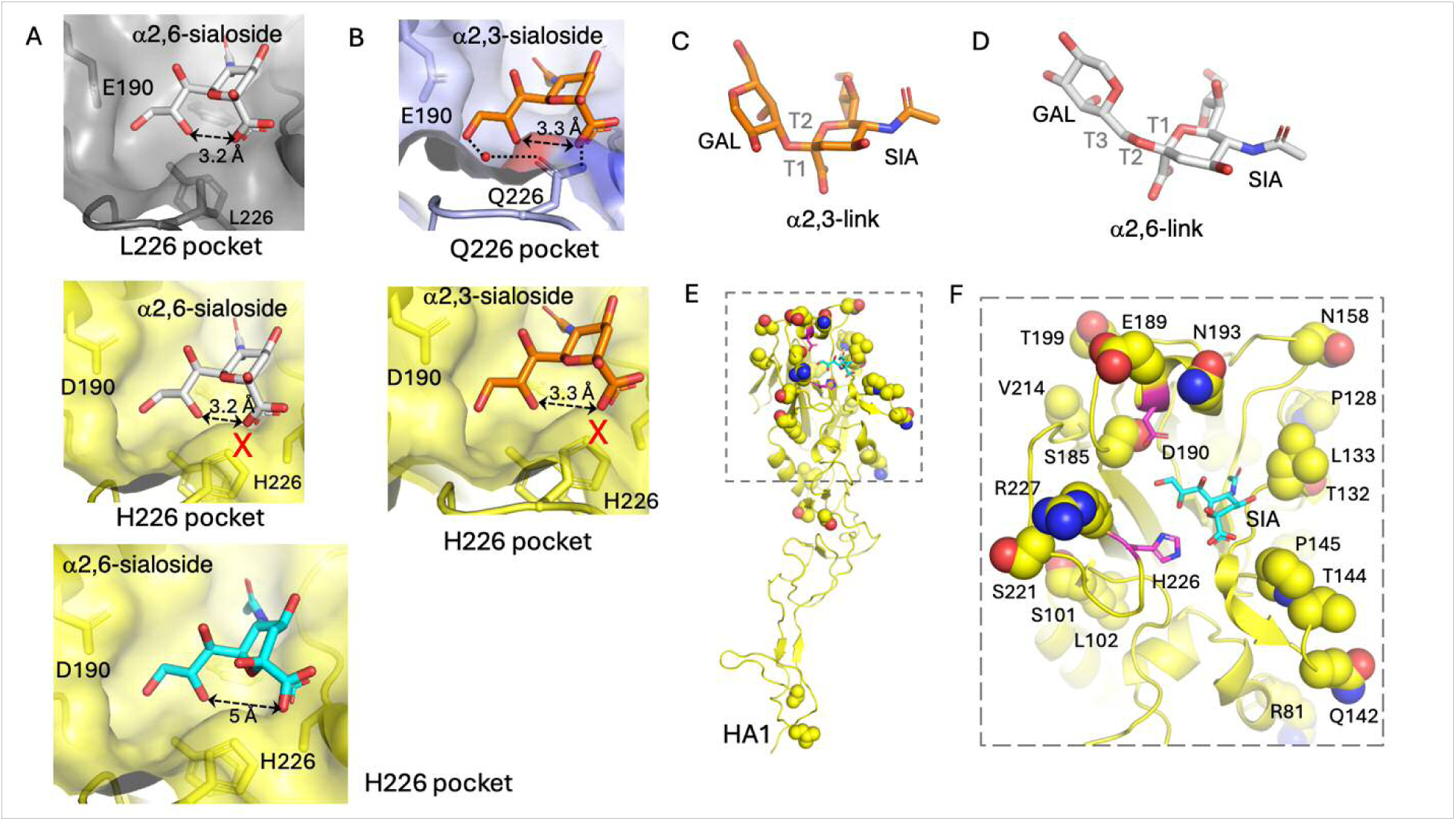
Binding of α2-6- vs α2-3-sialosides to the BC24-m HA pocket. **(A)** Binding of an α2-6-sialoside to L226 pocket (PDB 9DIO, top); this chair conformation of sialic acid is incompatible with H226 pocket of BC24-m HA (middle), and consequently, the sialic acid switches conformation to bind the mutant pocket (bottom). (**B**) The sialic acid of an α2-3-sialoside binds to BC24 HA with a canonical chair conformation (top); however, this conformation is not favored for binding to H226 pocket of BC24-m HA (bottom), explaining the reduced affinity of the mutant to α2-3-sialoside receptors. (**C**) An α2-3 linker has two rotatable bonds T1 and T2. (**D**) A higher torsional flexibility about T1, T2, and T3 bonds at the α2-6 link appears to help accommodate an α2-6-sialoside in H226 pocket. (**E**) Amino acid sequence comparison (Fig. S1B) of BC24 with TX24 or gGuan96 (A/goose/Guangdong/1/1996(H5N1)) HAs shows large sequence variations near the receptor binding site compared to other parts of HA. (**F**) A zoomed view of the site showing mutated residues relative to the other two H5N1 HAs; two mutated pocket residues, D190 and H226, are shown in magenta and sialic acid in cyan sticks. Mutations in this region may emerge to help in improving binding of α2-6-sialoside to BC24-m HA.

## Acknowledgements

The cryo-EM structures were determined at The Alberta CryoEM Facility and we would like to thank Dr. Zhijie Li for his help with the data collection. We would like to acknowledge Robert Smiley for bioinformatics support and Dr. Jack Moore at the Alberta Proteomics and Mass Spectrometry facility for mass spectrometry analysis. Molecular graphics and analyses performed with UCSF ChimeraX, developed by the Resource for Biocomputing, Visualization, and Informatics at the University of California, San Francisco, with support from National Institutes of Health R01-GM129325 and the Office of Cyber Infrastructure and Computational Biology, National Institute of Allergy and Infectious Diseases.

## Funding statement

Ministry of Technology and Innovation through Striving for Pandemic Preparedness—The Alberta Research Consortium (MG, KD, JSK)

Canada Excellence Research Chair Program (KD, LKM)

Alberta Innovates Graduate Student Scholarship (ZT)

Canada Biomedical Research Fund grant (APD)

Biosciences Research Infrastructure Fund grant (APD)

Natural Sciences and Engineering Research Council of Canada Discovery Grant (APD)

Natural Sciences and Engineering Research Council of Canada (JSK)

Canada Foundation for Innovation (JSK)

Alberta Innovation and Advanced Education Research Capacity Program (JSK)

## Author contributions (no punctuation in the initials)

Conceptualization: MG, KD, OFA

Methodology: MG, KD, JSK, APD, NZ, RAE, OFA, EPT, DK, ZT, DTB

Investigation: RAE, OFA, EPT, DK, EW, ZT, DTB, NZ

Visualization: MG, KD, JSK, APD, RAE, OFA, EPT, DK, EW, ZT, DTB

Funding acquisition: MG, KD, JSK, APD, LKM Project administration: MG, KD

Supervision: MG, KD, JSK, APD Writing – original draft: MG, KD, OFA

Writing – review & editing: MG, KD, JSK, APD, LKM, NZ, RAE, OFA, EPT, DK, EW, ZT, DTB

## Data, code, and materials availability

All data are available in the main text or the supplementary materials. The atomic coordinates have been deposited in the Protein Data Bank (PDB) under accession codes: 12QE; 12OL; 12QF; 12QG; 12RR; 12RS; 12RX; 12RW. The corresponding 3D electrostatic potential maps have been deposited in the Electron Microscopy Data Bank (EMDB) under accession codes: EMD-76680; EMD-76641; EMD-76681; EMD-76682; EMD-76714; EMD-76715; EMD-76720; EMD-76719.

## Supplementary Materials

## Other Supplementary Materials for this manuscript include the following

Validation reports for Structures I through VIII

## Materials and Methods

### Ethics statement

Identification and use of residual blood samples were carried out in accordance with local guidelines and regulations and approved by the University of Alberta Human Research Ethics Board (reference number Pro00150742).

### Cell lines and media

Human embryonic kidney (HEK) 293T cells were obtained from the American Type Culture Collection (ATCC) and maintained in D10 media (Dulbecco’s Modified Eagle Medium [DMEM; Gibco] supplemented with 10% heat-inactivated fetal bovine serum [FBS; Gibco]) at 37°C and 5% CO₂. Expi293F cells (Life Technologies), used for mammalian expression of recombinant Abs and hemagglutinin antigens, were maintained in Expi293F expression medium (Life Technologies) at 37°C and 8% CO₂. Insect *Sf9* cells were cultured at 27°C in *Sf9* SFM III Media (Invitrogen).

### Clinical samples

Specimens used in the study included combined 22 plasma and serum samples. These were residual samples collected from 22 individuals (age range of 4-73 years,) diagnosed with influenza via nucleic acid testing of respiratory specimens during the 2024/25 respiratory season who later had blood collected as part of their routine diagnostic care. Half of the participants were infected with influenza A H1H1 and half with H3N2. Collection dates for blood samples ranged from 15 to 180 days after the collection of the influenza-positive respiratory samples All participants resided in Alberta at the time of collection, a Canadian province with a population of 5.03 million (*47*). Immunization status and influenza infection in prior seasons were unknown.

### Generation of lentiviral pseudoviruses for serum neutralization assays

H5N1 pseudoparticles were generated according to methods adapted from (*29*). Briefly, 1 µg of a firefly luciferase–encoding lentiviral backbone (Addgene, #12260) was co-transfected into 50– 70% confluent HEK293T cells in a 6-well plate using TransIT-X2 transfection reagent, following the manufacturer’s recommendations, with 0.75 µg of Gag/Pol (Addgene, #170674), 0.1 µg of NA expression plasmid (BC24 NA), and 0.6 µg of H5 HA expression plasmid. Cell supernatants were collected 48 hours post-transfection, filtered through a 0.45 µm PES syringe filter (Millipore), and stored at −80°C until needed. H1N1 and H3N1 lentiviruses were generated similarly, but in 10 cm dishes (scaled appropriately) in the presence of 1.0 µg HAT (H1N1) or TMPRSS2 (H3N1). Clarified supernatants were concentrated over a 20% sucrose cushion in 1x PBS and ultracentrifuged for 2 hours at 4°C at 25,000 rpm using an SW28 rotor. Pellets containing concentrated particles were collected and stored at −80°C. Titers, reported as RLU/mL, were determined by serial dilution of virus stocks following methods described in (*29*), in the presence of 1 µg/mL TPCK-treated trypsin and 20 µg/mL polybrene for H1N1 and H3N1 pseudoviruses.

### Serum neutralization

For serum neutralization, 2e4 HEK293T cells were seeded per well in Poly-L-Lysine (Life Technologies) coated white-walled, 96-well plates (Life Technologies). ∼18hrs later, serially diluted sera incubated with 3e6RLU/mL of pseudovirus at 37°C for 1hr were transferred onto 293T cells. 48hr post infection, luciferase signal was measured using RLU as a proxy for infection count using the Bright-Glo Luciferase Kit (Promega). NT_50_ titers were determined from neutralization curves plotted with GraphPad (Prism 11).

### Expression and purification of recombinant hemagglutinins

The following hemagglutinin (HA) protein native sequences were downloaded from the Global Initiative on Sharing All Influenza Data (GISAID) (*48*): EPI176620|HA|A/California/07/2009|EPI_ISL_31158|A/H1N1(Cal09 HA); EPI2990337|HA|A/District Of Columbia/27/2023|EPI_ISL_18862356|A/H3N2 (Col23 HA); EPI3650014|HA|A/British_Columbia/PHL-2032/2024|EPI_ISL_19548836|A/H5N1 (BC24 HA), the sequence contains two X residues at positions 190 and 226 (H3 numbering) illustrating the amino acid heterogeneity; the combinations E190/Q226 and D190/H226 are referred to as “wild type” and “mutant” (BC24-m) hemagglutinin, respectively; EPI3171488|HA|A/Texas/37/2024|EPI_ISL_19027114|A/H5N1 (TX24 HA); EPI4800245|HA|A/Washington/2148/2025|EPI_ISL_20252012|A/H5N5 (WA25 HA); EPI1846961|HA|A/Astrakhan/3212/2020|EPI_ISL_1038924|A/H5N8 (Ast20 HA).

The ectodomain for each of the HAs was extracted from the native protein sequence and placed between the baculovirus gp67 secretion signal peptide (*49, 50*) (MVLVNQSHQGFNKEHTSKMVSAIVLYVLLAAAAHSAFAGS) at the N-terminus and the following amino acid sequence at the C-terminus (*51*): <u>GSGSGSPGS</u>*GYIPEAPRDGQAYVRKDGEWVLLSTFL*<u>GGSGGSGGS</u>**GRGVPHIVMVDAYK RYK**<u>GS</u>HHHHHHHH, where linkers are underlined, T4 fibritin foldon trimerization domain is italicized (*52*), Spy tag version 3 in bold letters (*53*), followed by an eight-histidine tag. Three previously reported trimer-stabilization substitutions (H355W, K380I, and E432I, H3 numbering) were introduced in the HA2 subunit (*54*). The H5N1 polybasic cleavage site was reduced to monobasic cleavage site to promote the stabilization of the HA in prefusion conformation. The corresponding coding DNA sequence was codon-optimized for expression in insect cells, synthesized and cloned into pBAC-1 (Invitrogen) by GenScript Biotech. The coding DNA sequence was transferred from pBAC-1 to linearized baculovirus DNA through recombination directly in *Sf9* insect cells using BestBac 2.0 Δ v-cath/chiA baculovirus co- transfection kit (Expression Systems). The recombinant baculovirus was amplified and used for expression of HA proteins according to the protocol by Drs. Garzoni, Bieniossek and Berger (*55, 56*). DNA sequence coding for the recombinant BC24 (E190/Q226) HA was also cloned into pcDNA3.4 plasmid for expression in in Expi293F cells (ATCC), except that the native secretion signal peptide was kept at the N-terminus of the HA ectodomain. The insect or human cell- expressed soluble HA was purified from the expression media though Ni-NTA (Thermo Fisher Scientific) affinity chromatography in phosphate-buffered saline (PBS), followed by size- exclusion chromatography (SEC) on Superose 6 increase 10/300 GL or Superdex 200 Increase 10/300 GL (Cytiva) column in PBS. For cryo-EM studies the purified insect-cell expressed HA was cleaved with trypsin at 1:50 ratio prior to SEC step (Promega). Purified proteins were flash- frozen on dry-ice methanol slurry and stored at -80⁰C until further usage.

### Ab expression, purification, and Fab preparation

Abs were expressed from human IgG1 and IgK expression vectors (Twist Biosciences) encoding the variable heavy and light genes, respectively, of selected Abs. Paired vectors were co- transfected at a 1:1 ratio using the Expi293F Expression System (Gibco) following the manufacturer’s protocol. 5-6 days post-transfection, culture media containing secreted Abs were clarified (2000 × g, 10 min), filtered, and passed through a Protein A GraviTrap column (Cytiva). Elution was carried out with 0.1 M glycine-HCl (pH 2.7), after which pooled Ab fractions were buffer-exchanged into 1x PBS using a 7 kDa MWCO Zeba desalting column (Life Technologies) and stored at −80°C until needed. Fabs were prepared by digesting purified IgGs at 37°C with immobilized papain (Life Technologies) according to the manufacturer’s protocol. Fabs were collected as the flow-through and first wash when digested fractions were passed through a rProtein A GraviTrap column (Cytiva). Fab-containing fractions were concentrated using a 10 kDa molecular weight cut-off spin column (Amicon), buffer-exchanged into 1× PBS, and stored at −80°C. IgG and Fab purity were assessed by SDS-PAGE.

### Hemagglutination inhibition assay

Serum samples were treated with receptor-destroying enzyme (RDEII, Denka Seiken Co.) overnight at 37°C, followed by heat inactivation at 56°C for 30 minutes min according to manufacturer instructions. Four hemagglutination units of purified hemagglutinin were combined with the serially two-fold diluted sera of the initial 1:10 dilution in U-shaped 96-well plates (Greiner), in a total volume of 15 µL and incubated at room temperature for 30 minutes. Subsequently, 15 µL of 1% turkey red blood cells (Rockland Inc.) was added and the plates that were incubated for 60-90 min at room temperature. HI titers were determined as the highest serum dilution that completely inhibited hemagglutination.

### Enzyme-linked immunosorbent assay (ELISA)

High-binding 96-well plates (Corning) were coated overnight at 4°C with 25 or 50 ng of hemagglutinins diluted in PBS, for monoclonal and pAbs, respectively. Plates were washed with PBS containing 0.05% Tween-20 (PBST) and blocked with 2% BSA (Roche) in PBST for 1 h at room temperature. After washing with PBST, wells were incubated with primary Abs (monoclonal or polyclonal), serially diluted in PBST with 0.1% BSA, for 1 h at room temperature. Wells were then washed with PBST and incubated with anti-IgG1 human secondary Abs conjugated with horseradish peroxidase (Invitrogen) for 1 h at room temperature. Plates were washed again and the signal was developed using tetramethylbenzidine substrate (Sigma Aldrich). The reaction was stopped with 2 M sulfuric acid after 30 min. Absorbance was measured at 450 nm using a microplate reader (SpectraMax M5, Molecular Devices). The signal, expressed as area under the curve (AUC), was calculated using Prism (GraphPad).

For the competitive ELISA, HA-coated (50 ng) and washed wells were pre-incubated with Fabs derived from 100F4 (4 µg/mL) and MEDI8852 (2 µg/mL) IgGs for 1 h at room temperature. Wells were then washed with PBST and the remaining steps were performed as described above. An HRP-conjugated secondary Ab (Invitrogen) recognizing Fc fragments was used to detect pAbs competing with the pre-bound Fabs.

### Immunoprecipitation and proteomic sample preparation for Immunoaffinity-mass spectrometry (IA-MS) analysis

High binding 96-well polystyrene microplates (Greiner Bio-One) were coated overnight with recombinant protein (0.3 µg/well), blocked with 5% sucrose in PBS, and washed, as previously reported(*57, 58*). Serum (20 μL) was diluted with 5% sucrose in PBS, incubated for two hours, and washed with 50 mM ammonium bicarbonate. Following two hours of on-plate incubation and washing, enriched Abs were reduced with dithiothreitol, alkylated with iodoacetamide, and digested on the same plate with trypsin (0.5 µg/well) (*59, 60*). Stable isotope-labelled absolute quantified peptide internal standards (SpikeTides_TQL) were synthesized, purified and quantified by JPT Peptide Technologies. Dithiothreitol, iodoacetamide, and MS-grade Trypsin were obtained from MilliporeSigma. SpikeTides_TQL peptide internal standards (300 fmol/well) were spiked before digestion(*61, 62*).

### Untargeted DIA LC-MS

Peptides were separated using the NanoElute2 system (Bruker, Billerica, MA) with a two-column separation mode with PepMap Neo Trap Cartridge (5 cm x 300 μm x 5 μm, 5 mm, #174500, Thermo Scientific) and PepSep analytical column (10 cm x 150 μm x 1.5 μm, Bruker) over 5-35% acetonitrile gradient for 7.5 min, at 600 nl/min flow rate and 40°C. Peptides were ionized with a Captive Spray 2 source and 20 µm emitter (Bruker) and analyzed with timsTOF Ultra 2 mass spectrometer (Bruker). Acquisition parameters included 200 °C source temperature, 1600 V voltage, positive mode, 3.0 L/min dry gas, and high sensitivity detection mode. Each dia-PASEF cycle (1.39 min) included one MS1 ramp (100 – 1700 m/z), 12 MS/MS ramps (331.3-1132.3 m/z) with 32 MS/MS windows over ≥+2 ion cloud (26 Da with 1 Da overlap), 0.65-1.30 1/K0 ion mobility range, and 100 ms ramp and accumulation times. Skyline software (v26.1) was used to process dia-PASEF files, extract the MS2 fragment peaks (MS2 filtering for DIA, centroid, all ions, 10 ppm) for the light endogenous peptides and heavy isotope-labelled SpikeTides_TQL peptide internal standards, and calculate the light-to-heavy (L/H) ratios. Serum volume (20 μL), internal standard amounts (300 fmol/well), and L/H ratios were used to calculate the absolute concentrations of each isotype and subclass of human Abs, as previously reported(*63*).

### Charge detection nMS

Charge detection (CD)-nMS measurements were performed on the UHMR mass spectrometer with the Direct Mass Technology (Thermo Fisher Scientific). The CD-MS data were processed using STORIBoard software (Proteinaceous).

### Native Mass Spectrometry High Throughput Screening

High-throughput screening of glycans binding to the HAs was performed using the Catch-and-Release (CaR)-nMS method (*38–40*). The HAs (1 µM) were mixed with glycan libraries (0.5-1 µM each). 2-5 µL of the solutions were loaded into the nano electrospray ionization (nanoESI) capillaries, which were produced from borosilicate glass capillaries (1.0 mm outside diameter (o.d.), 0.78 mm inner diameter (i.d.), 10 cm length) using a P-1000 micropipette (Sutter Instruments). CaR-nMS measurements were performed in negative mode using a Q Exactive Ultra-High Mass Range Orbitrap mass spectrometer (Thermo Fisher Scientific) equipped with a modified nanoflow ESI source. The nanoESI spray voltage was -0.7 to -0.8 kV. The capillary temperature was 150°C, and the S-lens RF level was 100; an automatic gain control target of 5×10^6^ and maximum injection time of 200 ms were used. The resolving power for collision-induced dissociation (CID) spectra was 25000, respectively. For the CaR-nMS measurements, HCD spectra were acquired using a range of collision energies (120-180 eV). Data acquisition and analysis were performed using Xcalibur version 4.4.

### Concentration Independent nMS

Quantitative measurements of HAs-glycan interactions were performed using the concentration - independent (COIN)-nMS measurements in CaR-nMS format as described previously (*41*). Briefly, two different solutions were loaded into the nanoESI capillaries. In both solutions, the HA of interest was prepared at the same concentration while the glycan concentration in solution 2 (30-50 µM) is higher than that of solution 1 (0.2-0.5 µM). Due to the concentration differences, mixing occurred over time leading to changes in the relative abundance of the complexes. To perform nanoESI, a voltage of approximately -0.7 kV was applied to a platinum wire inserted inside the nanoESI tip and in contact with the solution. The solution temperature was 25°C. Resolution (resolving power) of 25000 was used. Maximum injection time was 200 ms, the S-lens RF level was 200 and DC offset was 21. Collision energy was 120-160 V. Raw data were processed using the Thermo Xcalibur 4.4 software. Time-resolved mass spectra were averaged over 1 min intervals and the sum of the charge state-normalized abundances of the reactant and the complex ions were calculated automatically using the SWARM software (https://github.com/pkitov/CUPRA-SWARM) (*64*). The *K*_d_ values were fit using Igor pro (WaveMetrics Inc.) using eq 1:

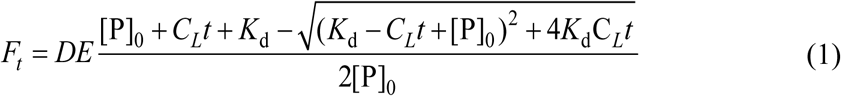

where DE is the detection efficiency of the released glycan (L) relative to the HAs (P), C_L_(*t*) is the time (*t*)-dependent function that describes the change in ligand concentration due to diffusion and advection, [P]_0_ is initial protein (HAs) concentration, the time-dependent fractional binding site occupancy (fraction bound, *F*_t_) of P, was calculated using the time-dependent abundance (*Ab*_t_) of the released ligand and free protein as shown in eq 2,

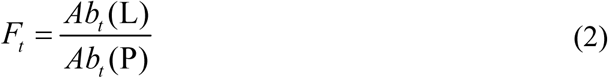

### Cryo-EM sample preparation

Media containing recombinantly expressed HA from insect *Sf9* or mammalian Expi293F cells were clarified by centrifugation at 1000× g for 20 minutes. The pH of the media was increased to pH 7.3 (at room temperature) by titrating with 1.0 M HEPES pH 7.7. A 4 mL bed volume of Ni- NTA resin (HisPur, Thermo Scientific) was washed in HA buffer (160 mM NaCl, 25 mM HEPES pH 7.3), added to the cleared supernatant and incubated for a minimum of two hours at 4°C. The media containing HA-bound Ni-NTA resin was poured over a ⌀2.5 cm x 20 cm Econo-Column (Bio-Rad), flow-through discarded, and resin washed four times with 25 mL of HA buffer. The washed resin was resuspended in a total volume of 14 mL HA buffer and transferred to a 15 mL conical tube. Trypsin (20 μg of sequencing grade modified trypsin #V5111, Promega) was resuspended in 0.5 mL of HA buffer, incubated for 30 minutes at room temperature, added to the HA-bound resin and incubated overnight on a tube rotator at 4°C. Proteolytic digestion processed HA0 into HA1/HA2 and cleaved HA from the resin-bound C-terminal trimerization domain, SpyTag (*65*) and octa-histidine tag. The digestion reaction was poured over an Econo-Column and the flow-through containing HA collected, concentrated to < 0.5 mL in an Amicon Ultra-4 centrifugal filter 30 kDa molecular weight cutoff device (MilliporeSigma) and further purified using size-exclusion chromatography (Superdex 200 Increase 10/300 GL, Cytiva). Peak fractions were pooled and concentrated.

### Cryo-EM Grid Preparation

Prior to grid preparation, detergent (n-Dodecyl-β-D-Maltoside; Life Technologies Inc #89902) was added to the sample to a final concentration of 0.005% w/v. Sample supports (Quantifoil Micro Tools GmbH Carbon R 1.2/1.3 on copper 200 or UltrAuFoil Gold R 1.2/1.3 on gold 300) were glow discharged in air using a Pelco easiGlow cleaning system for 45 s at −25 mA (copper grids) or 100 s at −15 mA (gold grids). A Thermo Fisher Scientific Inc Vitrobot Mark IV system was used to blot and vitrify 3 uL of sample by plunge freezing into liquid ethane. Instrument parameters were set as follows: chamber humidity 100%; chamber temperature 4°C; blot time 3–4 s; blot force 1; wait time 30 s.

### Cryo-EM Data Collection and Image Processing

Micrographs were collected at nominal magnifications of either 130000× or 165000× (raw pixel sizes 0.94 Å/pixel or 0.74 Å/pixel, respectively) on a 300 kV transmission electron microscope Krios G4 equipped with a X-FEG field emission gun, Selectris X energy filter and Falcon 4i direct electron detector. The movies were recorded in either EER or TIFF format. Total electron dose for each exposure was in the range 41–45 e^-^/Å^2^, under-focus targets were set in the EPU preset between -1 and -2.0, at steps of -0.2 μm. All image processing was performed in cryoSPARC (*66*) (version v4.7.1-cuda12). While volume reconstruction workflows varied by sample, they generally followed the undermentioned procedures (cryoSPARC native algorithms unless otherwise stated). Patch motion correction and CTF estimation followed by initial particle picking using cryoSPARC’s blob picking implementation. Two rounds of 2D classification and removal of junk classes was followed by a three-class ab initio reconstruction to form an initial assessment of heterogeneity. Subsequent 3D refinement of a selected ‘good’ class. A random subset of particles from the ‘good’ class was used as a positive training set to train a Topaz (*67*) (version 0.25.a) model. Topaz picking using the trained model produced a basis particle set. Further 2D and 3D multiclass decoy refinement produced a single consensus particle set and volume. The consensus particle set were regenerated using reference-based motion correction. A procedure was developed to ascertain, and deconvolute if present, heterogenous Fab occupancy of the head domain. The appropriate trimeric hemagglutinin-Fab model was first docked into the consensus volume using ChimeraX (*68*). Models of the eight possible Fab binding permutations of the HA-Fab complex (fully occupied; no Fab binding, three each of singularly and doubly occupied) were created and the molmap command used to simulate volumes of each model at 8 Å resolution. The volumes were imported into cryoSPARC and used as starting volumes for 3D classification of the consensus particle set. In samples where the molmap classification procedure cleanly separated the partially bound states, the subset of particles representing the fully bound HA-Fab complex was used as the basis for final non-uniform and local refinement with C3 symmetry enforced. The potential heterogeneous occupancy of the stem-binding Fab MEDI8852 was ignored due to its primary role as a stabilizing domain in the complex, and the increased number of permutations (64) that would need to be tested, leading to particle sets too small to produce high-resolution 3D volumes.

### Model Building and Refinement

Initial models were docked into the cryo-EM density map using ChimeraX and energy minimized using the ISOLDE (*69*) plugin. Pair-wise trimeric distance and dihedral restraints were applied during minimization. Models were further built using COOT (*70*) with final refinement using PHENIX (*71*) real-space refinement. Model statistics, including Ramachandran plots, clashscores, and rotamer outliers, were analyzed using MolProbity (*72*) to ensure optimal geometric quality and are reported in Supplemental Table S3.

### Phylogenetic analysis

For tree construction, the selected Influenza A strains were approved for use in vaccines by the FDA in the United States and EMA in the European Union according to an FDA briefing in 2024 (*73*). The strain A/American_Wigeon/South_Carolina/22-000345-001/2021, which was approved for vaccine use in Canada (*74*), was also selected for the phylogenetic tree inference. In addition, all remaining (> 800) H5 sequences in the GISAID database, which originated human hosts, were clustered using CD-hit (*75*) with the similarity threshold of 0.95 resulting in 16 representative sequences which were used for tree construction as well. Finally, the HA from A/Korea/426/1968(H2N2) was chosen as an outgroup for later rooting of the tree. Thus, selected sequences were aligned using MAFFT (*76, 77*) using the --localpair and –maxiterate 1000 options. The resulting alignment was then run through an automated trimming process using TrimAl (*78*), with the -automated1 option. Phylogenetic inference was performed using PhyML (*79*) with the following options: --search=SPR --datatype=aa --model=WAG. Finally, the resulting tree was rendered using the Interactive Tree of Life web software (*80*), and edited using Inkscape (The Inkscape Project).

## Supplementary Figures

**Fig. S1.**
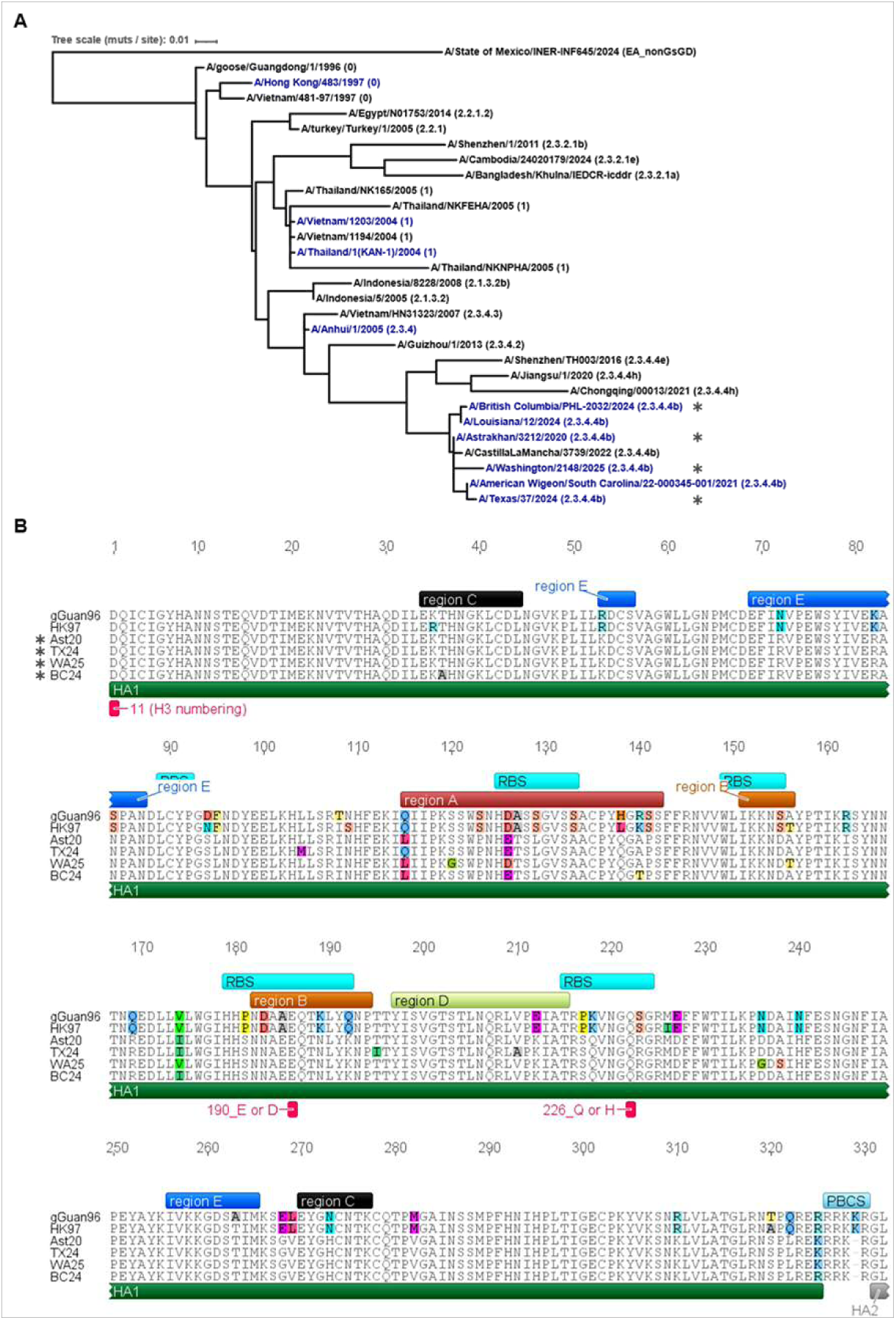
Phylogenetic relationships between a selected set of H5 hemagglutinin sequences. Asterisks indicate the hemagglutinins used in this study. (**A**) Phylogenetic tree of H5 hemagglutinin sequences from the viruses targeted in Canada and the U.S. by licensed vaccines (blue font), as well as example H5 sequences from the indicated clades. The tree was generated using CD-Hit, MAFFT, TrimAl, PhyML, and Interactive Tree of Life (ITOL). Branch lengths illustrate the average number of mutations per site relative to the node that they are connected to. The scale for branch lengths is indicated. (**B**) Sequence alignment of a selected set of hemagglutinins (HA1 domain): gGuan96, A/goose/Guangdong/1/1996(H5N1); HK97, A/Hong Kong/483/1997(H5N1), Ast20, A/Astrakhan/3212/2020(H5N8); TX24, A/Texas/37/2024(H5N1); WA25, A/Washington/2148/2025(H5N5); BC24, A/BC/PHL-2032/2024(H5N1). Numbers above and below (in red) the sequences indicate residue numbers according to the H5 and H3 numbering, respectively. The locations of the RBS and the antigenic regions are according to Dadonaite et al., 2024 (*46*). PBCS – polybasic cleavage site, RBS – receptor binding site. The alignment and the image were generated using Geneious 5.5 software.

**Fig. S2.**
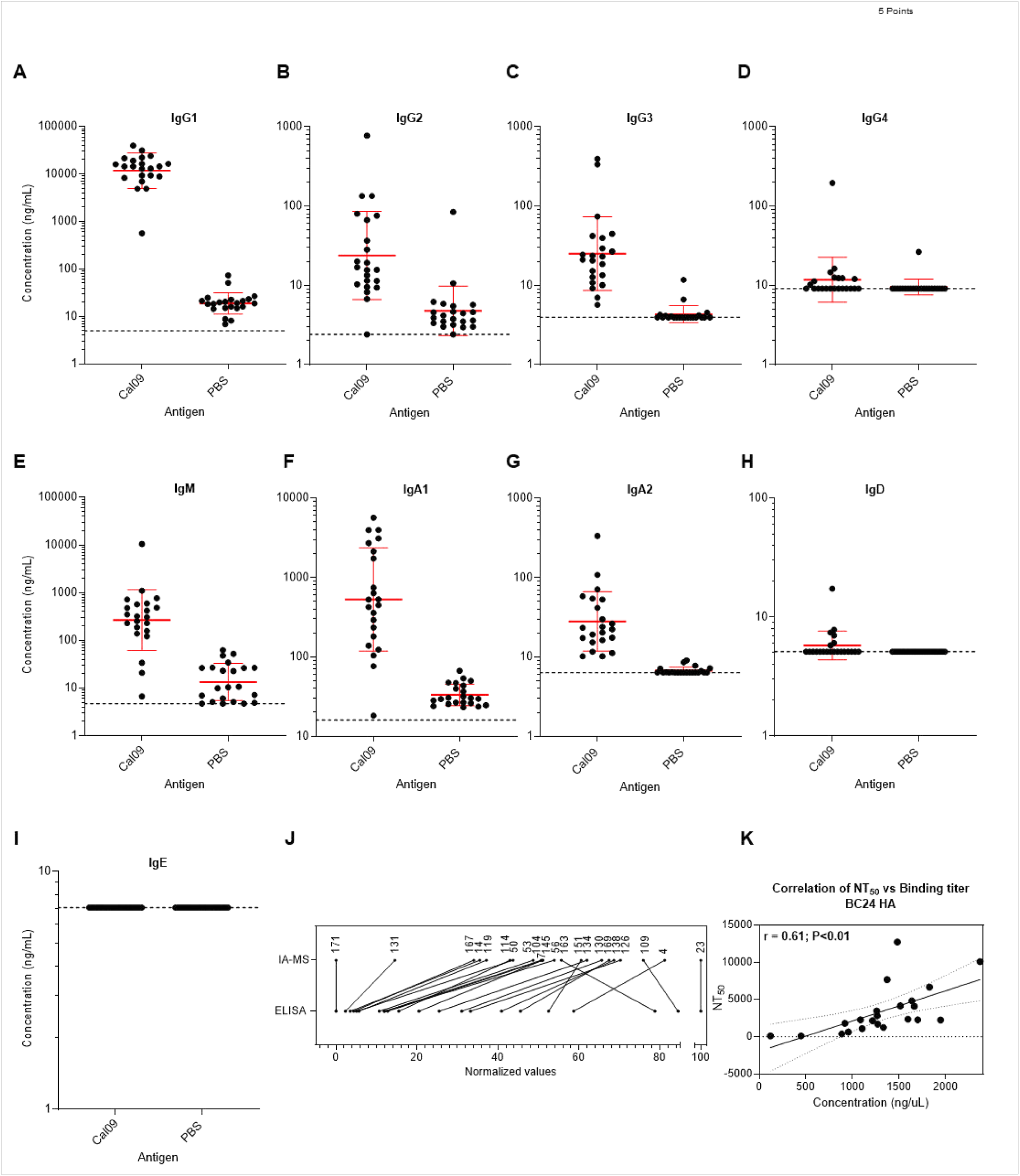
Serum immunoglobulin profiling and assay correlations. **(A-I)** Absolute concentrations of serum/plasma immunoglobulin (Ig) isotypes binding to recombinant Cal09 HA and PBS-negative control, quantified by immunoaffinity mass spectrometry (IA-MS). Values are shown as ng/mL; error bars indicate geometric mean concentration ± SD factor. Dotted lines indicate calculated limit of detection (see Table S2). A-I; as labelled on each graph (**J**) Correlation between normalized IgG1 measurements obtained by IA-MS and ELISA-derived IgG1 area under the curve (AUC) values from the same samples. Each point (connected) is an individual sample (identifier indicated). (**K**) Correlation between serum IgG1 PBS-subtracted absolute concentrations (x-axis) and pseudovirus neutralization titers (NT₅₀; y-axis) against BC24 HA. r, Pearson correlation coefficient.

**Fig. S3.**
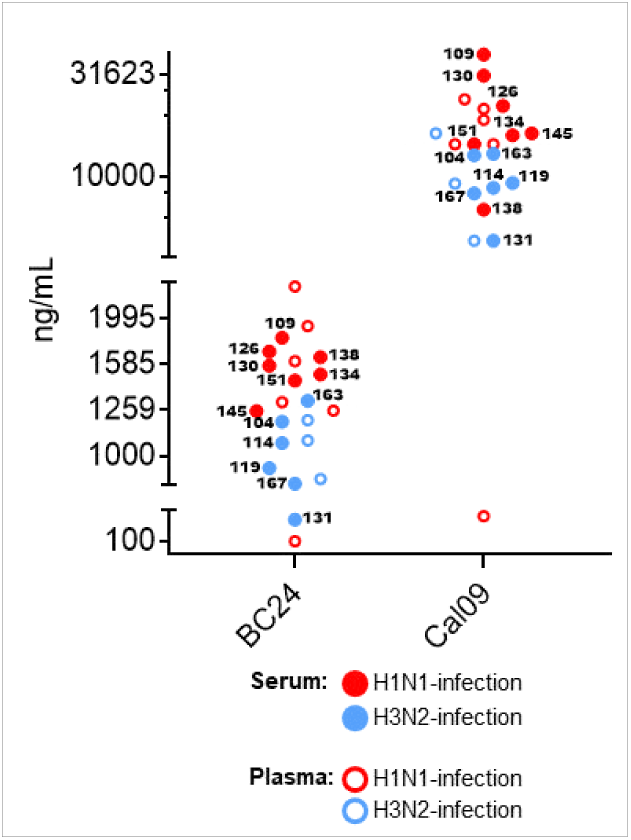
Hemagglutination inhibition titers for serum samples and IgG1 concentration in serum and plasma detected by IA-MS. Closed circles (labeled with sample IDs) represent sera analyzed by HI and MS, the open circles depict plasma samples analyzed solely by MS. Color coding for clinical diagnosis: red, H1N1 infection; blue, H3N2 infection. The serum subset of clinical samples used for HI was representative of H1N1- and H3N2-infected individuals and spanned the IgG1 range for both BC24 and Cal09 HAs.

**Fig. S4.**
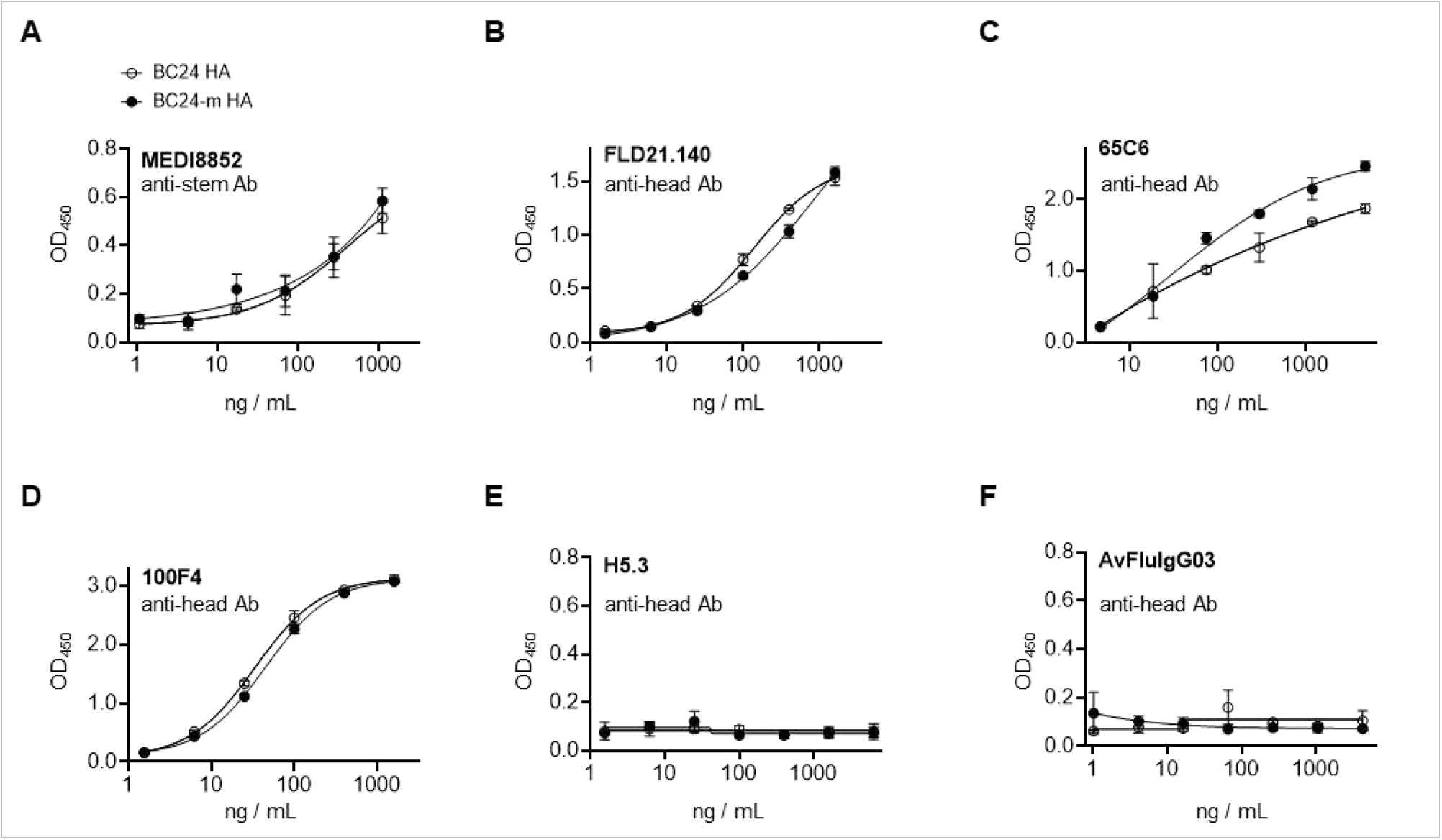
Ab binding to BC24 and BC24-m HAs, quantified by ELISA. Five anti-H5N1 HA mAbs targeting the head domain along with the stem-binding Ab MEDI8852, were analysed: (**A**) MEDI8852; (**B**) FLD21.140; (**C**) 65C6; (**D**) 100F4; (**E)** H5.3; and (**F**) AvFluIgG03. The HAs were digested with trypsin.

**Fig. S5.**
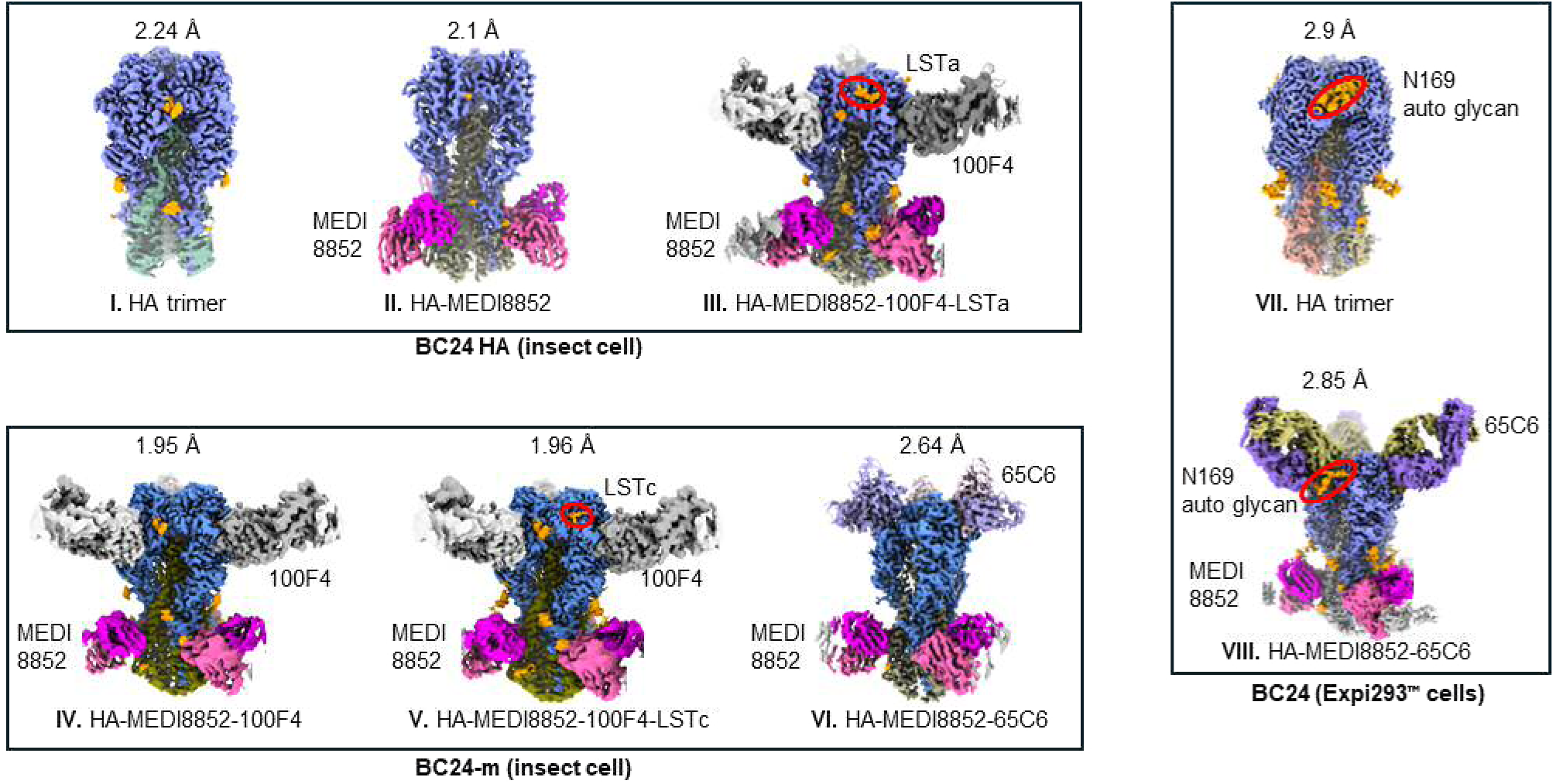
Structures of BC24 and BC24-m complexes. Eight structures of apo and various complexes of HA with Abs (MEDI8852, 100F4, and 65C6) and avian-receiptor-like glycan LSTa or human-receiptor-like glycan LSTc or an N169 glycosylated auto-glycan. The structures of insect-cell expressed BC24 and BC24-m HAs are grouped in top- and bottom-left boxes respectively. Expi293F-expressed BC24 HA structures are in the right box. The experimental cryo-EM density maps are colored as: trimeric HAs (blue), MEDI8852 (magenta/pink), 100F4 (light/dark gray), 65C6 (purple/olive), glycans and auto- glycans (orange).

**Fig. S6.**
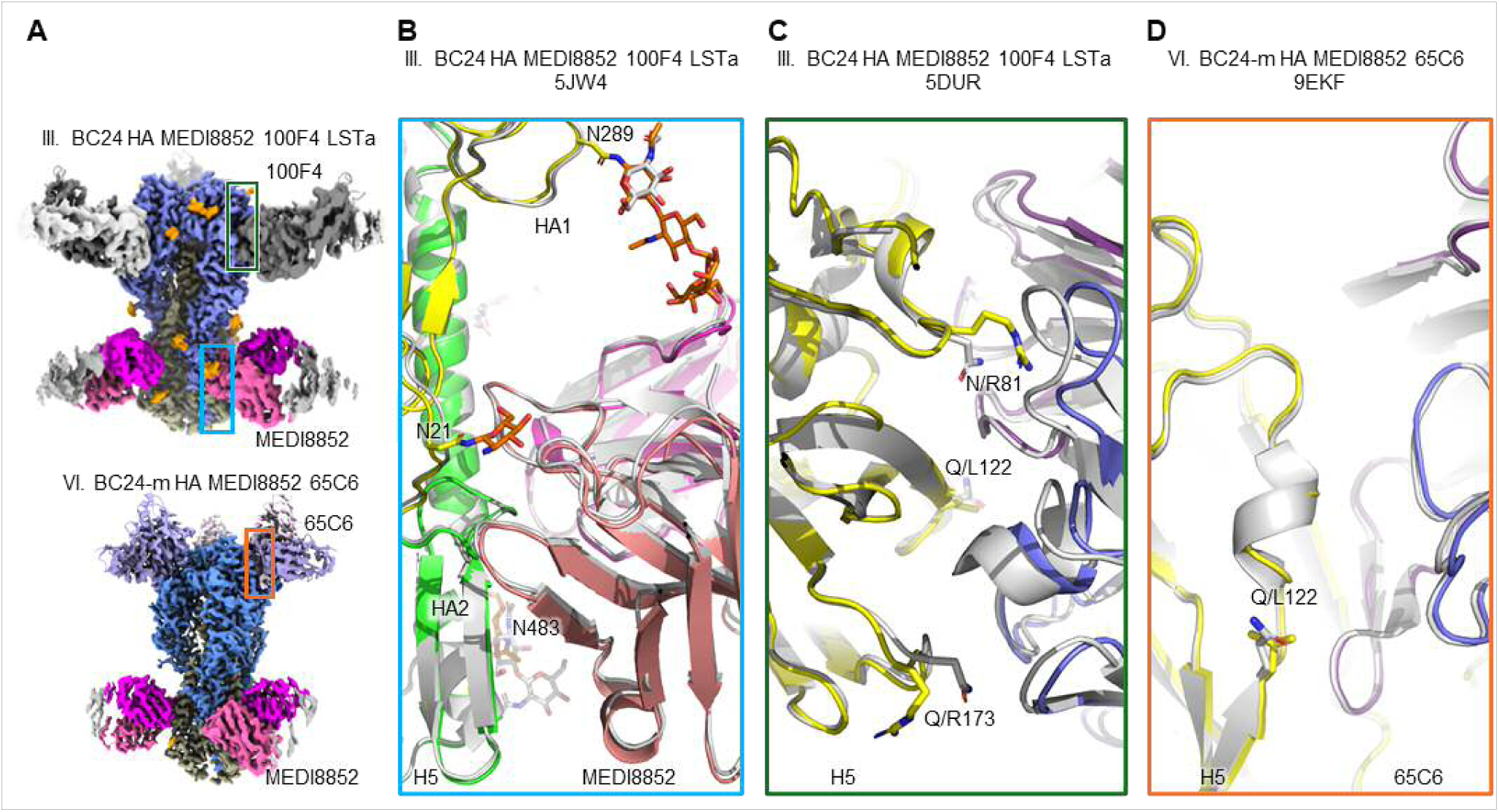
Epitope interface of three mAbs bound to BC24 HA. (**A**) Overview of BC24/ MEDI8852/100F4/LSTa structure (top) and BC24-m HA/MEDI8852/65C6 structure (bottom). The boxed areas indicate the close-ups shown in B, C and D panels. (**B**) Interacting region of the stem-binding mAb MEDI8852 (salmon/pink) with BC24 HA (yellow HA1, green HA2) is compared to that in 5JW4 (gray); the auto-glycans are in orange. Asn289 and Asn21 N-linked glycan from BC24 HA are shown interacting with the MEDI8852. (**C**) Head-binding mAb 100F4 (blue/purple) interacting with BC24 HA (yellow) is superimposed on an 100F4 complex (PDB: 5DUR, gray). The HA amino acid differences at the interface repositions 100F4. (D) Head-binding mAb 65C6 (blue/purple) interacting with BC24-m HA (yellow) is superimposed on TX24 HA-65C6 complex (PDB: 9EFK, gray). The mAb-HA interactions are conserved in two superimposed structures. The structures were aligned on HA Cα atom superpositions.

**Fig. S7.**
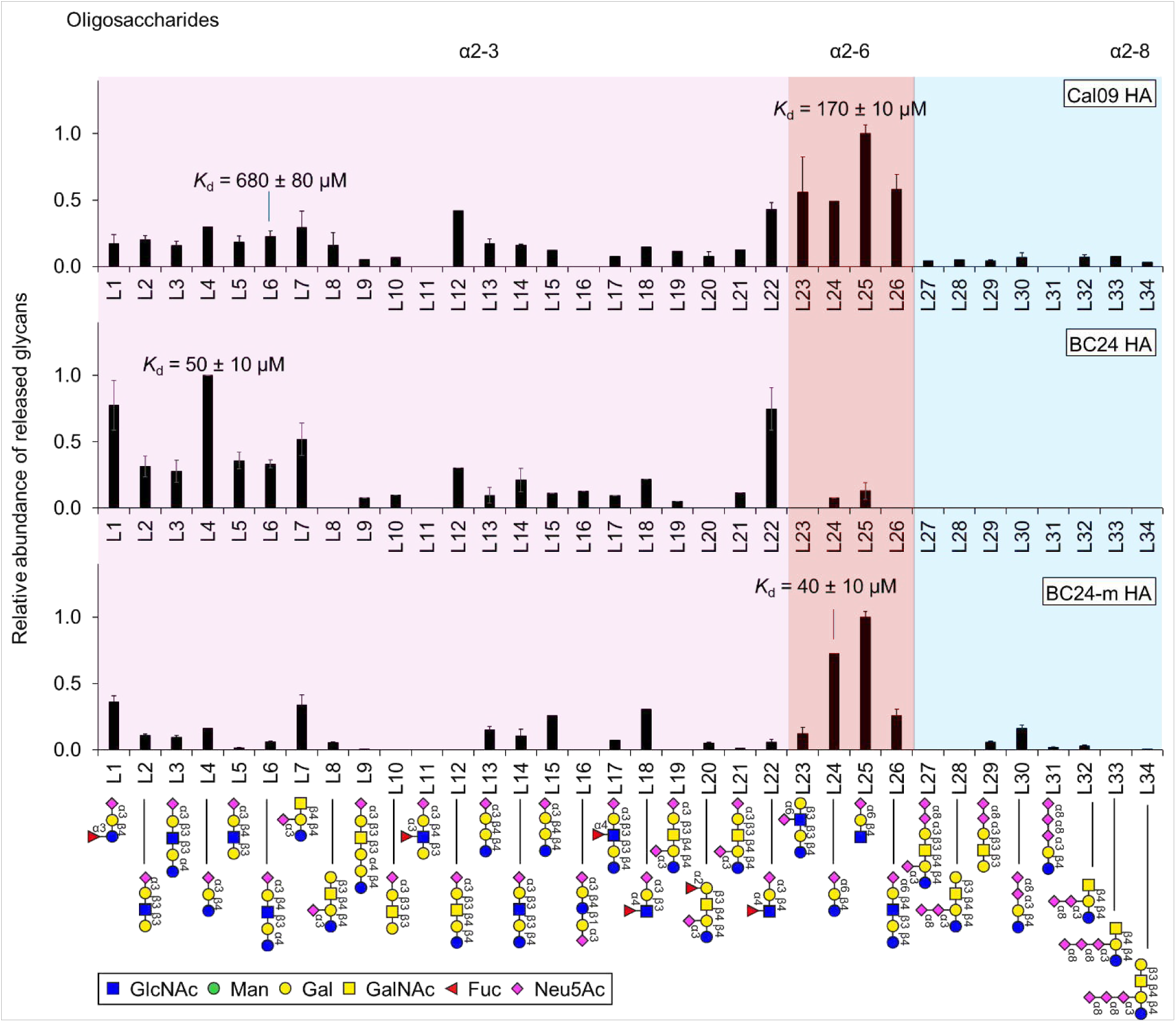
Relative abundance of released oligosaccharides from Cal09, BC24 and BC24-m HAs. The glycan binding was obtained from the Catch-and-Release native mass spectrometry (CaR-nMS) experiments and the *K*_d_ values obtained from the Concentration Independent (COIN)-nMS measurements. Measurements were performed as described in Methods. Data represent mean ± standard deviation obtained from 4 replicates. The glycans are presented using the symbol nomenclature for glycans (SNFG).

**Fig. S8.**
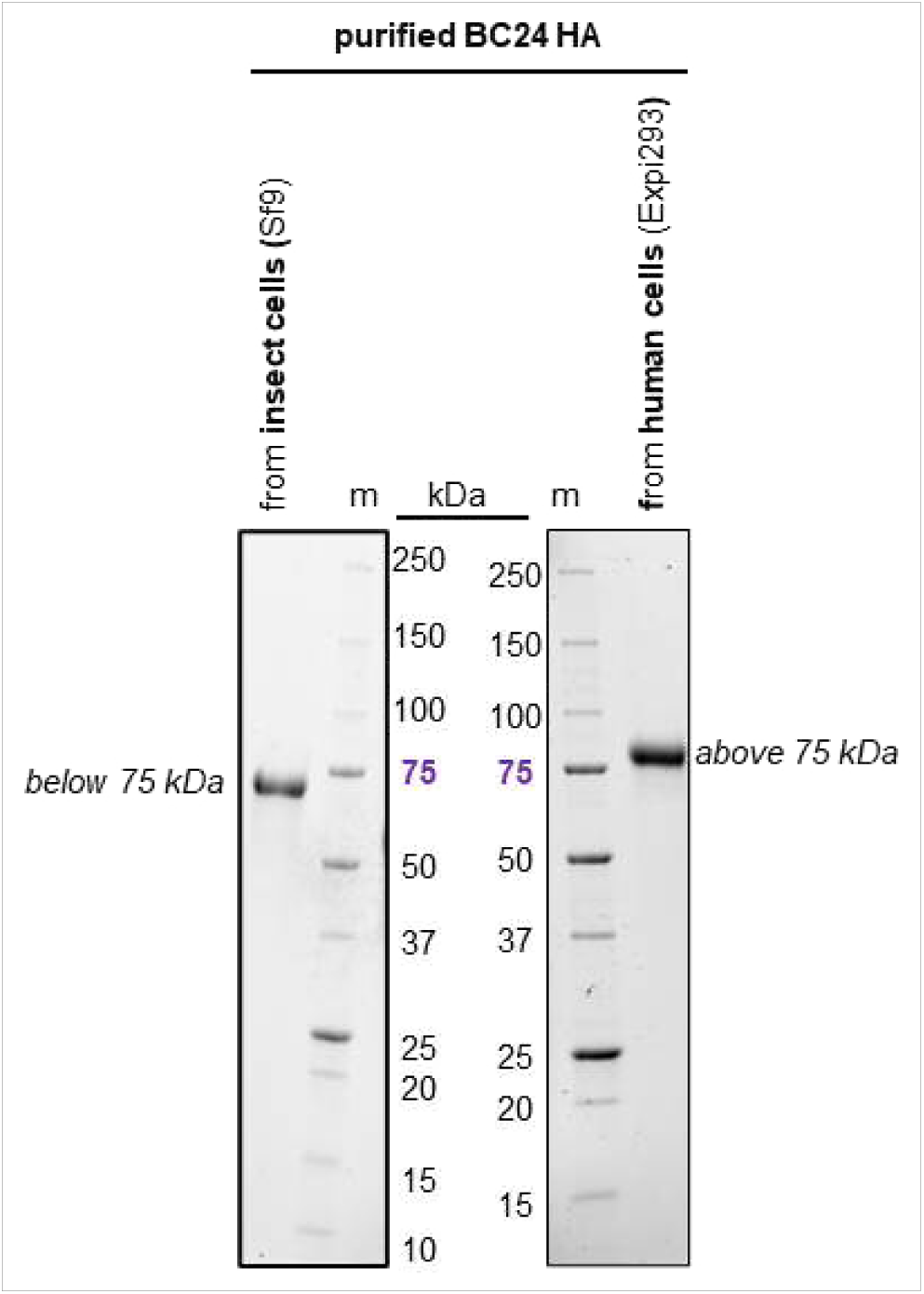
Denaturing PAGE migration patterns of BC24 HA expressed and purified from insect (left) versus human cells (right). The molecular weight markers show that the *Sf9* cell- expressed HA is below and Expi293 cell-expressed HA is above 75 kDa marker.

**Fig. S9.**
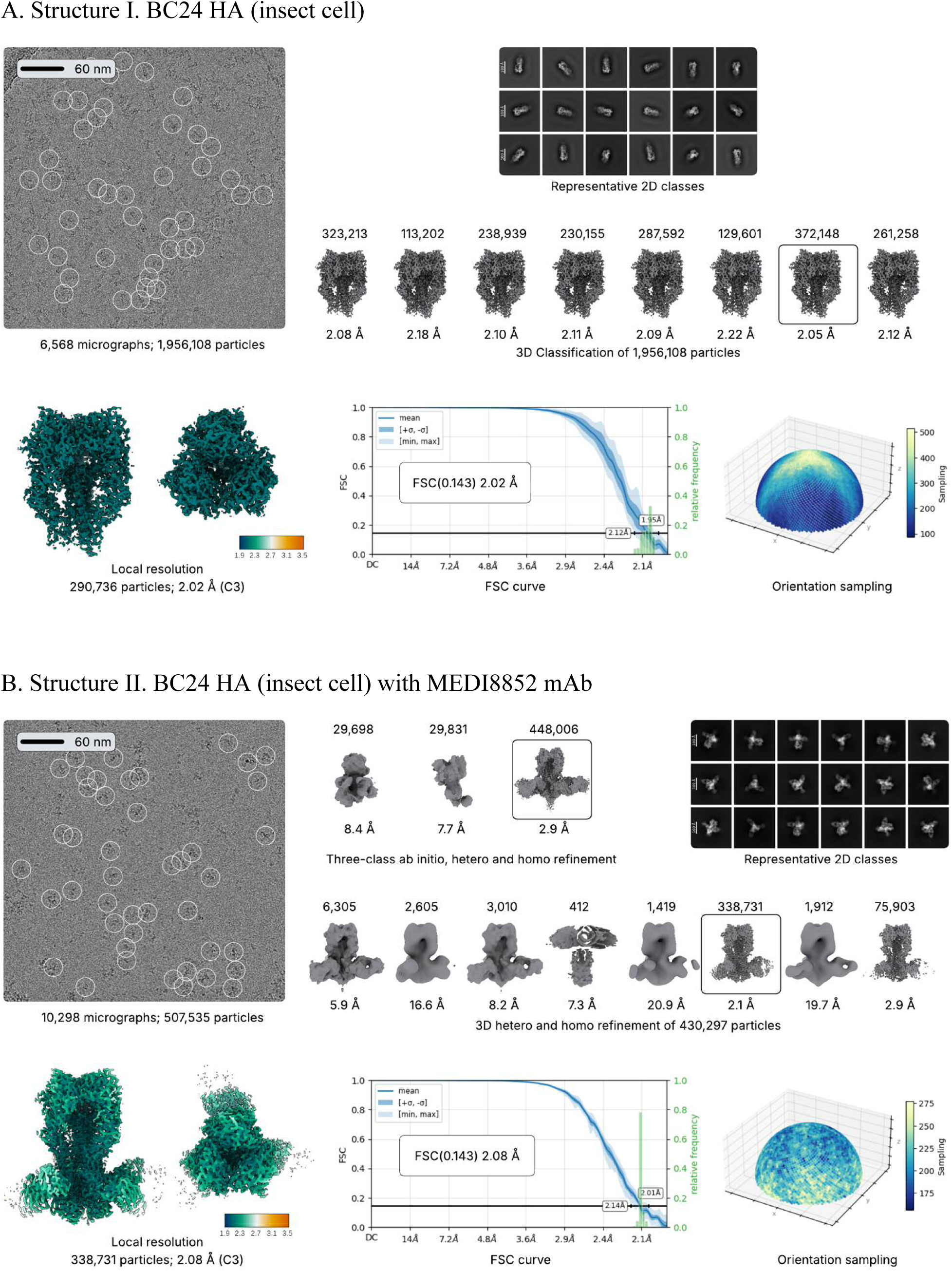
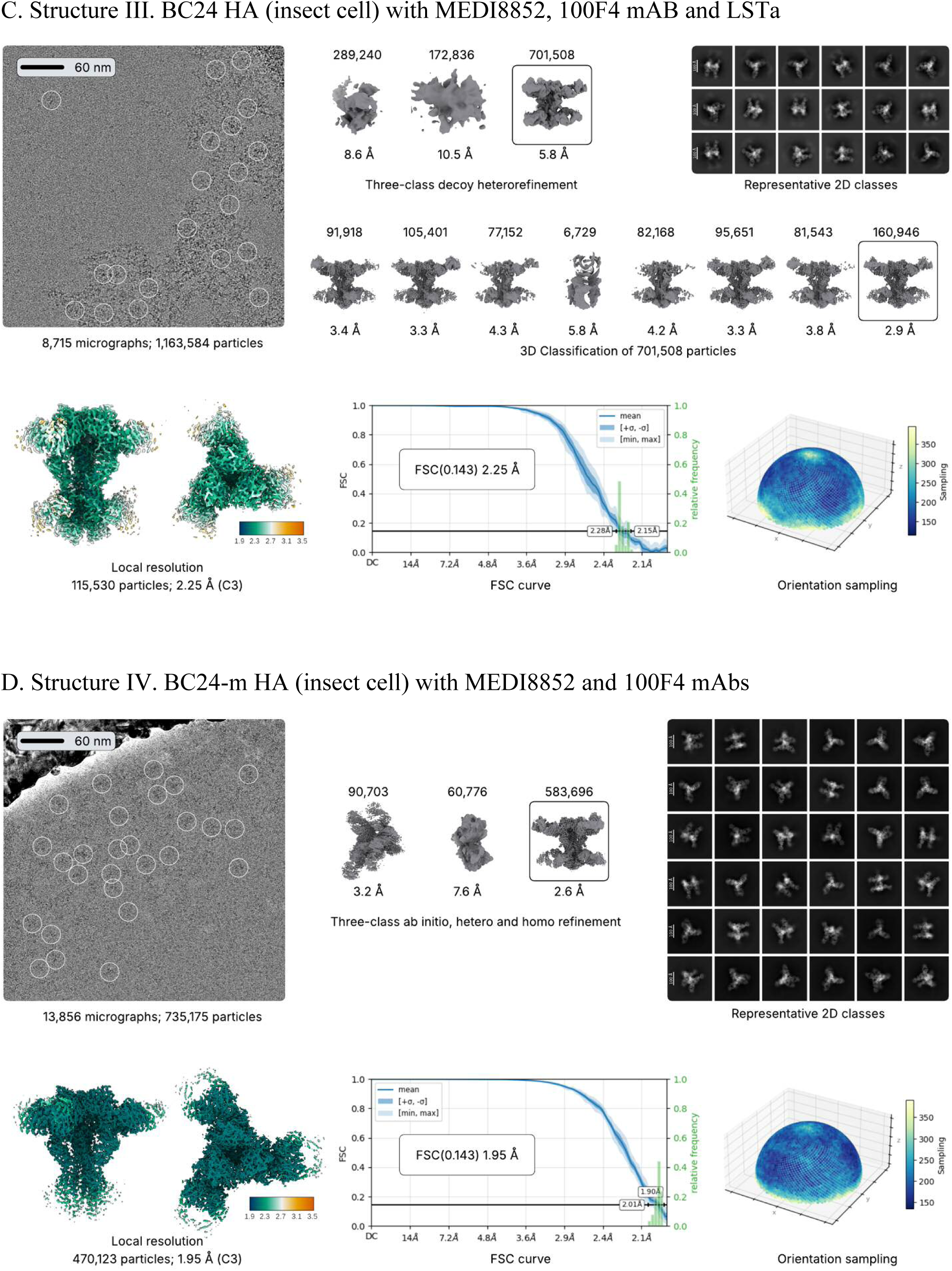
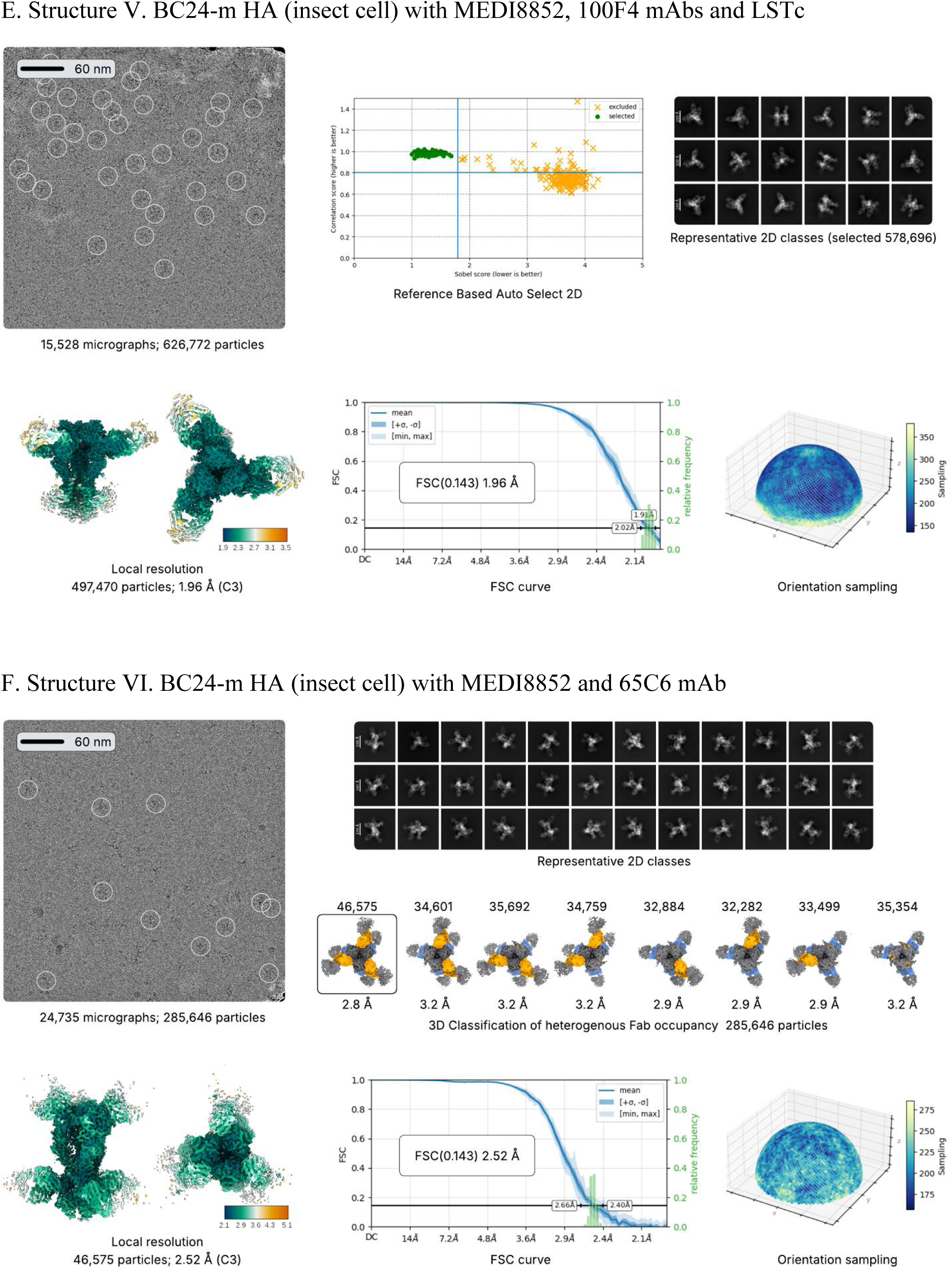
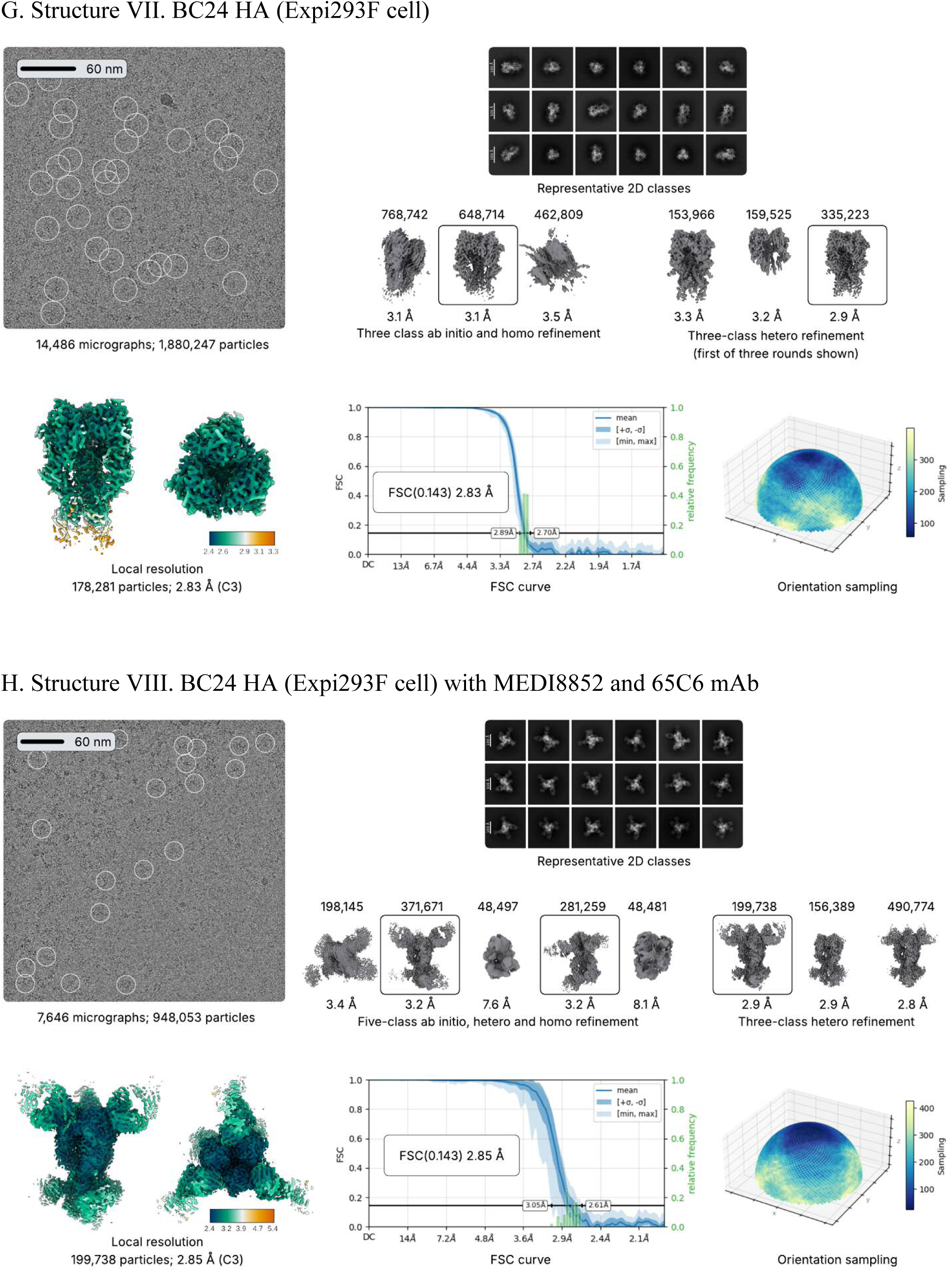
CryoEM processing and assessment. **(A-H)** Representative micrograph with 2 Å low-pass filter applied; the final particles are circled. A representative set of selected 2D classes from 2D classification of picked particles. The 3D volumes generated from a cleaned set of particles are shown with the number of particles used and the map resolution. The particle set constituting the boxed volume was carried forward for generating the final density map. The final cryo-EM density map for each complex was calculated using C3 symmetry, and each map is shown in color by local resolution; the attached color bar represents the resolution range. The gold-standard Fourier shell correlation (FSC) curve of the final reconstruction is based on the FSC = 0.143 criterion. Particle orientation distribution is shown by mapping Fourier sampling on half of a spherical surface. The detailed data processing parameters are in Table S3.

## Supplementary tables

**Table S1.** Participants characteristics summary in this study Group Study population.

| Group | Study population<br>(2024-2025 Flu season) |
| --- | --- |
| Number of participants | 22 |
| Age in years (median/Range) | 49 (4-73) |
| Sex |  |
| Influenza infection status |  |
| H1N1 Positive | 12 (55.5%) |
| H3N2 Positive | 10 (45.5%) |
| Blood sample collection time <sup>a</sup> |  |
| Early (< 40 days) | 13 (59%) |
| Late (> 40 days) | 9 (41%) |
| Influenza vaccination prior to infection | Unknown |
| Comorbidities | Unknown |

<sup>a</sup> Sample collection date: March to June 2025

**Table S2.** Serum concentrations of anti-Cal09 immunoglobulin isotypes and subclasses, as measured by IA-MS. A total of 22 PCR confirmed positive H1N1 or H3N2 clinical samples were tested with (+) Cal09 H1 antigen and (–) PBS. To calculate statistical significance, concentrations below LOD were adjusted to the LOD levels.

|  | IgG1 | IgG2 | IgG3 | IgG4 | IgM | IgA1 | IgA2 | IgD | IgE |
| --- | --- | --- | --- | --- | --- | --- | --- | --- | --- |
|  | ng/mL<sup>1</sup> |  |  |  |  |  |  |  |  |
| (+) H1N1 | 14415 | 17 | 22 | 9 | 314 | 486 | 23 | 5.1 | 7.1 |
| [IQR]<sup>2</sup> | [8,690-19,636] | [10-71] | [12-40] | [9.1-12.3] | [153-575] | [170-2,268] | [16-53] | [5.1-6.0] | [7.1-7.1] |
| [GMC±SD]<sup>3</sup> | **11729±**<br>**2** | **24±4** | **25±3** | **12±2** | **266±4** | **526±4** | **23±2** | **6±1** | **7±1** |
| (–) PBS | 19 | 4 | 4.0 | 9.1 | 11 | 29 | 6.4 | 5.1 | 7.1 |
| [IQR] | [15.1-22.7] | [3.3-5.5] | [4.0-4.2] | [9.1-9.1] | [5.1-27] | [26-44] | [6.4-7.0] | [5.1-5.1] | [7.1-7.1] |
| [GMC±SD] | 19±2 | 5±2 | 4±1 | 10±1 | 13±3 | 33±1 | 7±1 | 5±1 | 7±1 |
| Fold change | 759 | 4 | 6 | 1 | 29 | 17 | 4 | 1 | 1 |
| *W*<sup>4</sup> *P*-value | 4.8e-7 | 7.5e-7 | 4.8e-7 | 9.0e-2 | 4.7e-7 | 9.5e-7 | 4.7e-7 | 3e-2 | n/a<sup>5</sup> |

<sup>1</sup> median concentrations, <sup>2</sup> [IQR], interquartile ranges, <sup>3</sup> [GMC±SD], geometric mean concentrations and SD factor, <sup>4</sup> *W*, a 2-tail Wilcoxon matched-pairs signed rank test, <sup>5</sup> n/a, cannot calculate a paired test.

**Table S3.**
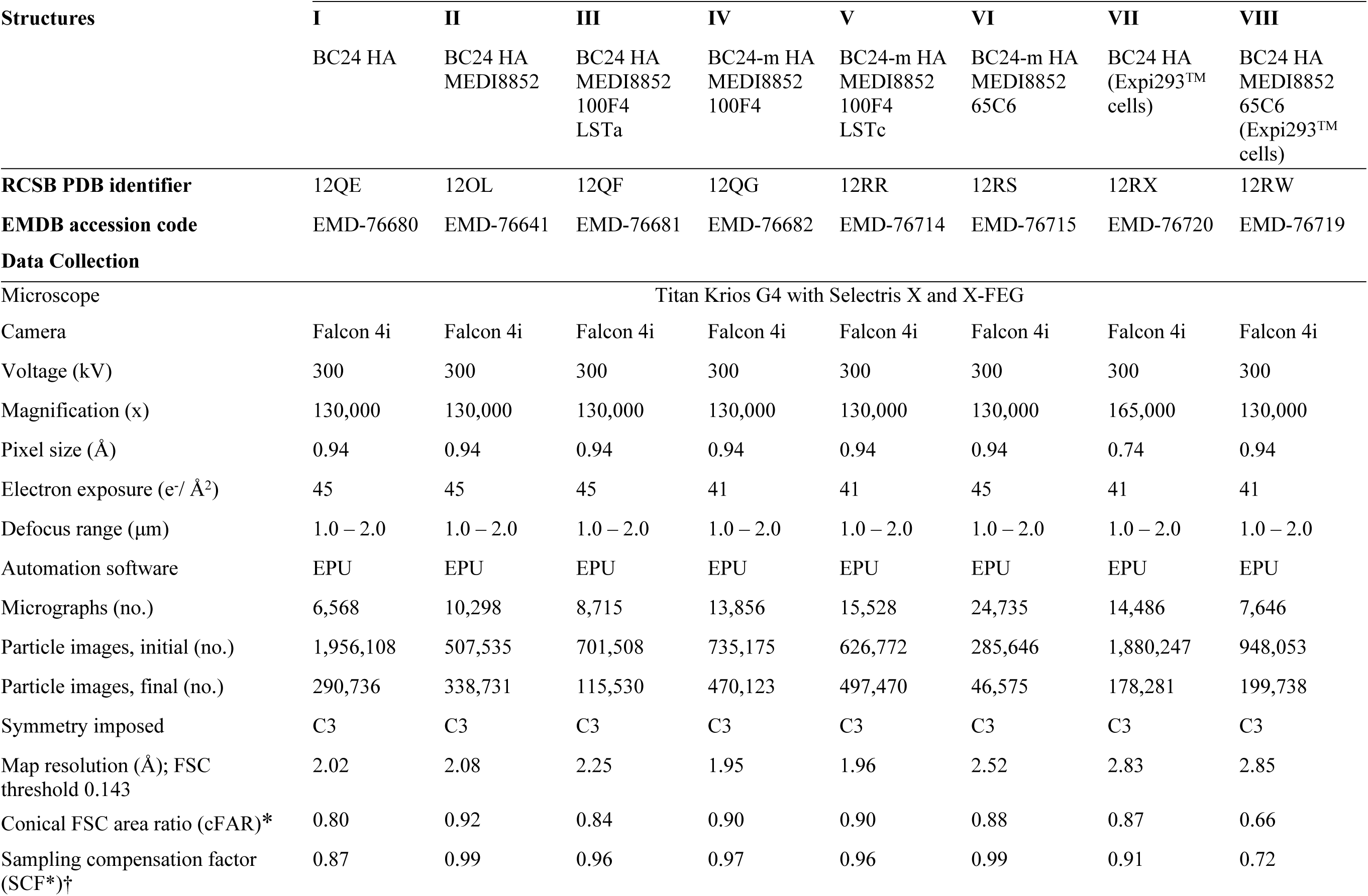
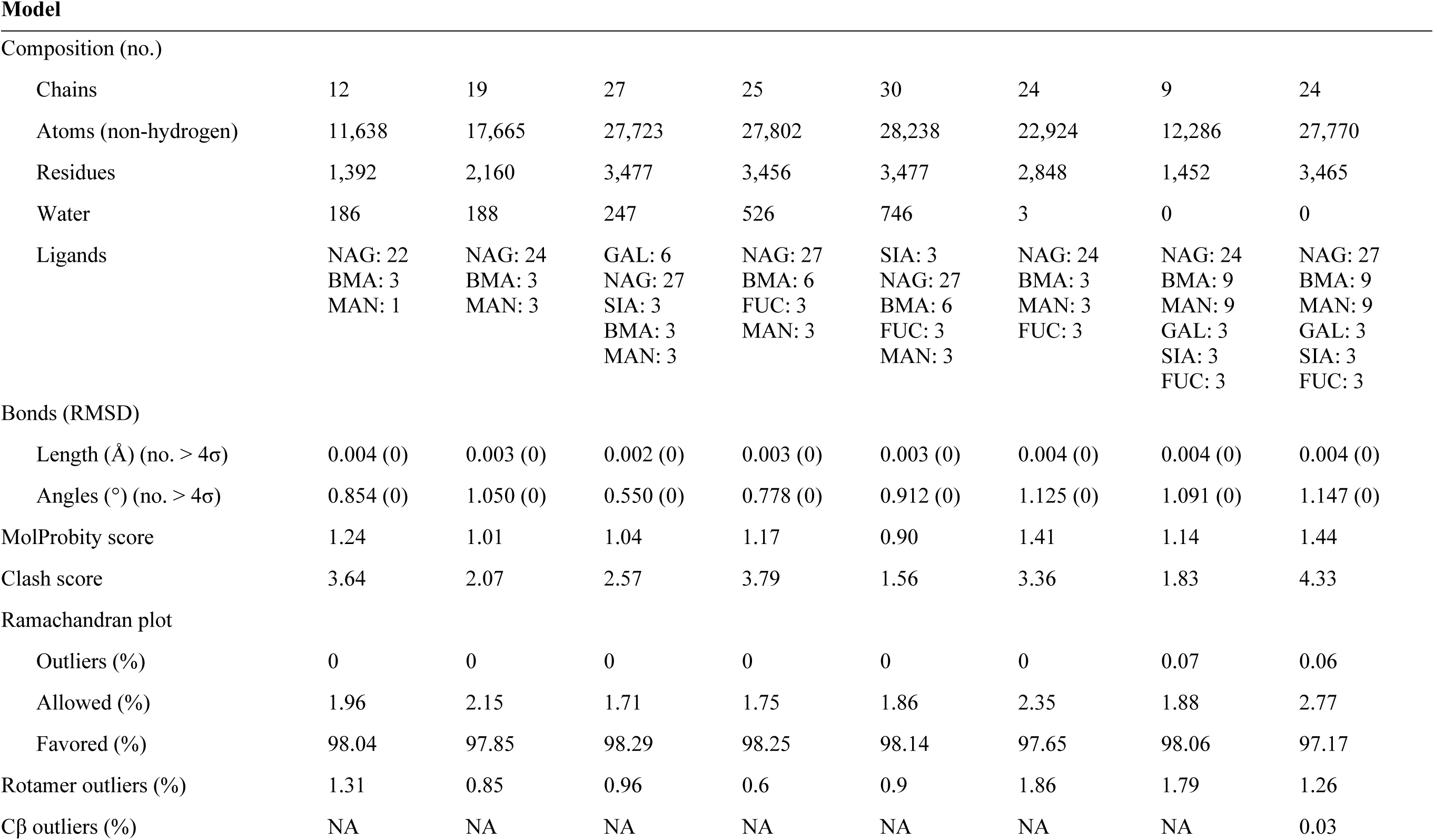
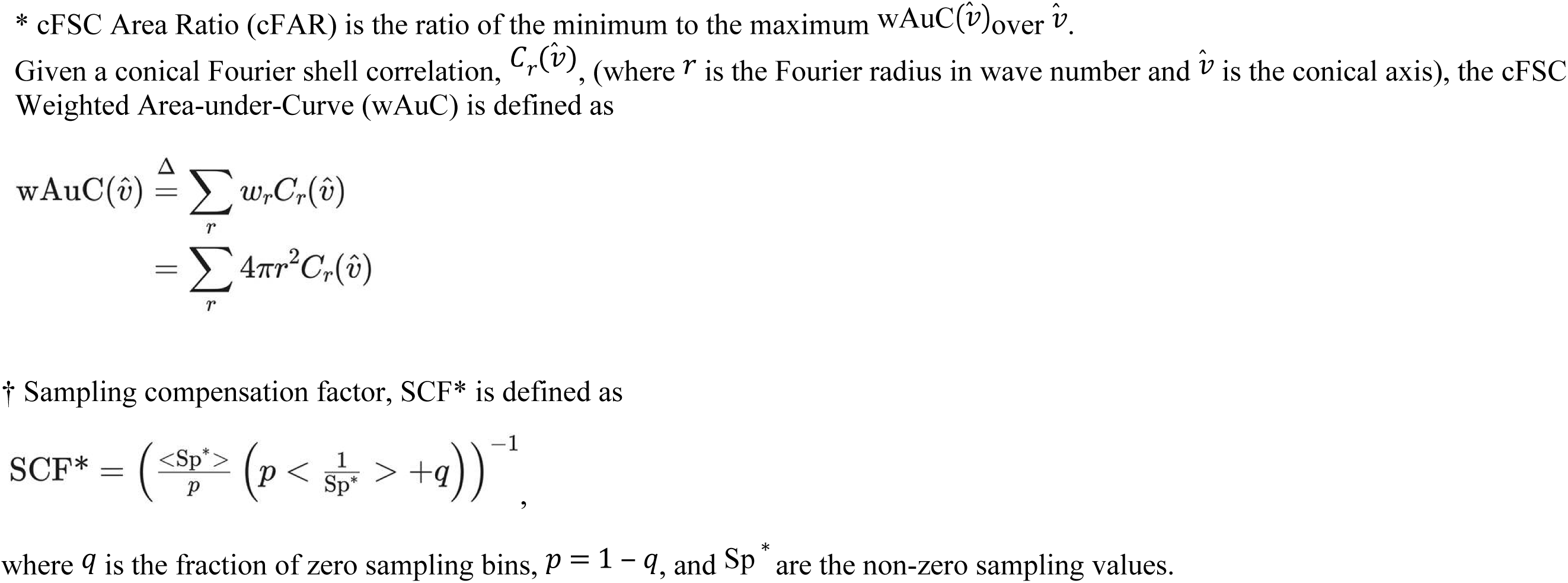
CryoEM Data Collection and Model Validation Statistics.

| **Structures** | **I** | **II** | **III** | **IV** | **V** | **VI** | **VII** | **VIII** |
| --- | --- | --- | --- | --- | --- | --- | --- | --- |
|  | BC24 HA | BC24 HA<br>MEDI8852 | BC24 HA<br>MEDI8852<br>100F4<br>LSTa | BC24-m HA<br>MEDI8852<br>100F4 | BC24-m HA<br>MEDI8852<br>100F4<br>LSTc | BC24-m HA<br>MEDI8852<br>65C6 | BC24 HA<br>(Expi293<sup>TM</sup><br>cells) | BC24 HA<br>MEDI8852<br>65C6<br>(Expi293<sup>TM</sup><br>cells) |
| **RCSB PDB identifier** | 12QE | 12OL | 12QF | 12QG | 12RR | 12RS | 12RX | 12RW |
| **EMDB accession code** | EMD-76680 | EMD-76641 | EMD-76681 | EMD-76682 | EMD-76714 | EMD-76715 | EMD-76720 | EMD-76719 |
| **Data Collection** |  |  |  |  |  |  |  |  |
| Microscope | Titan Krios G4 with Selectris X and X-FEG |  |  |  |  |  |  |  |
| Camera | Falcon 4i | Falcon 4i | Falcon 4i | Falcon 4i | Falcon 4i | Falcon 4i | Falcon 4i | Falcon 4i |
| Voltage (kV) | 300 | 300 | 300 | 300 | 300 | 300 | 300 | 300 |
| Magnification (x) | 130,000 | 130,000 | 130,000 | 130,000 | 130,000 | 130,000 | 165,000 | 130,000 |
| Pixel size (Å) | 0.94 | 0.94 | 0.94 | 0.94 | 0.94 | 0.94 | 0.74 | 0.94 |
| Electron exposure (e<sup>-</sup>/ Å<sup>2</sup>) | 45 | 45 | 45 | 41 | 41 | 45 | 41 | 41 |
| Defocus range (µm) | 1.0 – 2.0 | 1.0 – 2.0 | 1.0 – 2.0 | 1.0 – 2.0 | 1.0 – 2.0 | 1.0 – 2.0 | 1.0 – 2.0 | 1.0 – 2.0 |
| Automation software | EPU | EPU | EPU | EPU | EPU | EPU | EPU | EPU |
| Micrographs (no.) | 6,568 | 10,298 | 8,715 | 13,856 | 15,528 | 24,735 | 14,486 | 7,646 |
| Particle images, initial (no.) | 1,956,108 | 507,535 | 701,508 | 735,175 | 626,772 | 285,646 | 1,880,247 | 948,053 |
| Particle images, final (no.) | 290,736 | 338,731 | 115,530 | 470,123 | 497,470 | 46,575 | 178,281 | 199,738 |
| Symmetry imposed | C3 | C3 | C3 | C3 | C3 | C3 | C3 | C3 |
| Map resolution (Å); FSC threshold 0.143 | 2.02 | 2.08 | 2.25 | 1.95 | 1.96 | 2.52 | 2.83 | 2.85 |
| Conical FSC area ratio (cFAR)* | 0.80 | 0.92 | 0.84 | 0.90 | 0.90 | 0.88 | 0.87 | 0.66 |
| Sampling compensation factor (SCF*)† | 0.87 | 0.99 | 0.96 | 0.97 | 0.96 | 0.99 | 0.91 | 0.72 |
| **Composition (no.)** |  |  |  |  |  |  |  |  |
| Chains | 12 | 19 | 27 | 25 | 30 | 24 | 9 | 24 |
| Atoms (non-hydrogen) | 11,638 | 17,665 | 27,723 | 27,802 | 28,238 | 22,924 | 12,286 | 27,770 |
| Residues | 1,392 | 2,160 | 3,477 | 3,456 | 3,477 | 2,848 | 1,452 | 3,465 |
| Water | 186 | 188 | 247 | 526 | 746 | 3 | 0 | 0 |
| Ligands | NAG: 22<br>BMA: 3<br>MAN: 1 | NAG: 24<br>BMA: 3<br>MAN: 3 | GAL: 6<br>NAG: 27<br>SIA: 3<br>BMA: 3<br>MAN: 3 | NAG: 27<br>BMA: 6<br>FUC: 3<br>MAN: 3 | SIA: 3<br>NAG: 27<br>BMA: 6<br>FUC: 3<br>MAN: 3 | NAG: 24<br>BMA: 3<br>MAN: 3<br>FUC: 3 | NAG: 24<br>BMA: 9<br>MAN: 9<br>GAL: 3<br>SIA: 3<br>FUC: 3 | NAG: 27<br>BMA: 9<br>MAN: 9<br>GAL: 3<br>SIA: 3<br>FUC: 3 |
| **Bonds (RMSD)** |  |  |  |  |  |  |  |  |
| Length (Å) (no. > 4σ) | 0.004 (0) | 0.003 (0) | 0.002 (0) | 0.003 (0) | 0.003 (0) | 0.004 (0) | 0.004 (0) | 0.004 (0) |
| Angles (°) (no. > 4σ) | 0.854 (0) | 1.050 (0) | 0.550 (0) | 0.778 (0) | 0.912 (0) | 1.125 (0) | 1.091 (0) | 1.147 (0) |
| MolProbity score | 1.24 | 1.01 | 1.04 | 1.17 | 0.90 | 1.41 | 1.14 | 1.44 |
| Clash score | 3.64 | 2.07 | 2.57 | 3.79 | 1.56 | 3.36 | 1.83 | 4.33 |
| **Ramachandran plot** |  |  |  |  |  |  |  |  |
| Outliers (%) | 0 | 0 | 0 | 0 | 0 | 0 | 0.07 | 0.06 |
| Allowed (%) | 1.96 | 2.15 | 1.71 | 1.75 | 1.86 | 2.35 | 1.88 | 2.77 |
| Favored (%) | 98.04 | 97.85 | 98.29 | 98.25 | 98.14 | 97.65 | 98.06 | 97.17 |
| Rotamer outliers (%) | 1.31 | 0.85 | 0.96 | 0.6 | 0.9 | 1.86 | 1.79 | 1.26 |
| Cβ outliers (%) | NA | NA | NA | NA | NA | NA | NA | 0.03 |

\* cFSC Area Ratio (cFAR) is the ratio of the minimum to the maximum $\text{wAuC}(\hat{v})_{\text{over } \hat{v}}$ .
Given a conical Fourier shell correlation, $C_r(\hat{v})$ , (where $r$ is the Fourier radius in wave number and $\hat{v}$ is the conical axis), the cFSC Weighted Area-under-Curve (wAuC) is defined as
† Sampling compensation factor, SCF\* is defined as
where $q$ is the fraction of zero sampling bins, $p = 1 - q$ , and $\text{Sp}^*$ are the non-zero sampling values.

## References and Notes

1. R. M. Azeem et al., Emerging threats of H5N1 clade 2.3.4.4b: cross-species transmission, pathogenesis, and pandemic risk. Front Cell Infect Microbiol 15, 1625665 (2025).

2. A. N. Jassem et al., Critical Illness in an Adolescent with Influenza A(H5N1) Virus Infection. New Engl J Med 392, (2025).

3. N. M. C. De Jong, A. Aartse, M. J. Van Gils, D. Eggink, Development of broadly reactive influenza vaccines by targeting the conserved regions of the hemagglutinin stem and head domains. Expert Rev Vaccines 19, 563–577 (2020).

4. T. M. Uyeki et al., Highly Pathogenic Avian Influenza A(H5N1) Virus Infection in a Dairy Farm Worker. New Engl J Med 390, 2028–2030 (2024).

5. U.S. Centers for Disease Control and Prevention. (U.S. Centers for Disease Control and Prevention, U.S. Centers for Disease Control and Prevention, 2024).

6. U.S. Centers for Disease Control and Prevention. (U.S. Centers for Disease Control and Prevention, U.S. Centers for Disease Control and Prevention, 2024).

7. C. M. Sheppard et al., An Influenza A virus can evolve to use human ANP32E through altering polymerase dimerization. Nat Commun 14, (2023).

8. E. Staller et al., Structures of H5N1 influenza polymerase with ANP32B reveal mechanisms of genome replication and host adaptation. Nat Commun 15, (2024).

9. Z. H. Zhu, H. T. Fan, E. Fodor, Defining the minimal components of the influenza A virus replication machinery via an in vitro reconstitution system. Plos Biol 21, (2023).

10. J. S. Long, B. Mistry, S. M. Haslam, W. S. Barclay, Host and viral determinants of influenza A virus species specificity. Nature Reviews Microbiology 17, 67–81 (2019).

11. J. J. S. Santos et al., Bovine H5N1 binds poorly to human-type sialic acid receptors. Nature 640, E18–E20 (2025).

12. P. Chopra et al., Receptor-binding specificity of a bovine influenza A virus. Nature 640, E21–E27 (2025).

13. A. J. Eisfeld et al., Pathogenicity and transmissibility of bovine H5N1 influenza virus. Nature 633, 426–432 (2024).

14. J. H. Ni et al., Loss of α2,3-linked sialoside in the receptor-binding site of a H5N1 influenza hemagglutinin identified in a human patient. bioRxiv, 2026.2001.2019.700419 (2026).

15. E. Kovács, M. Ríos Carrasco, M. F. Guerreiro Cabana, R. P. de Vries, H5N1 2.3.4.4b HA E190D and Q226H mutations, picked up as minority variants in a patient, result in an inability to bind sialic acid. bioRxiv, 2026.2003.2006.710037 (2026).

16. S. Wilks, M. de Graaf, D. J. Smith, D. F. Burke, A review of influenza haemagglutinin receptor binding as it relates to pandemic properties. Vaccine 30, 4369–4376 (2012).

17. J. Stevens et al., Structure and receptor specificity of the hemagglutinin from an H5N1 influenza virus. Science 312, 404–410 (2006).

18. G. N. Rogers et al., Single amino acid substitutions in influenza haemagglutinin change receptor binding specificity. Nature 304, 76–78 (1983).

19. T. H. Lin et al., A single mutation in bovine influenza H5N1 hemagglutinin switches specificity to human receptors. Science 386, 1128–1134 (2024).

20. T. M. Tumpey et al., A two-amino acid change in the hemagglutinin of the 1918 influenza virus abolishes transmission. Science 315, 655–659 (2007).

21. B. F. Koel et al., Substitutions near the receptor binding site determine major antigenic change during influenza virus evolution. Science 342, 976–979 (2013).

22. C. Gioia et al., Cross-subtype immunity against avian influenza in persons recently vaccinated for influenza. Emerg Infect Dis 14, 121–128 (2008).

23. N. C. Morano et al., Structure of a zoonotic H5N1 hemagglutinin reveals a receptor- binding site occupied by an auto-glycan. Structure 33, 228–233 e223 (2025).

24. Z. N. Li et al., Pre-existing cross-reactive immunity to highly pathogenic avian influenza 2.3.4.4b A(H5N1) virus in the United States. Nat Commun 16, 10954 (2025).

25. V. Le Sage et al., Pre-existing H1N1 immunity reduces severe disease with bovine H5N1 influenza virus. bioRxiv, (2024).

26. K. H. Restori et al., Preexisting immunity to the 2009 pandemic H1N1 virus reduces susceptibility to H5N1 infection and disease in ferrets. Sci Transl Med 17, eadw4856 (2025).

27. Y. Rais, A. P. Drabovich, Identification and Quantification of Autoantibodies against Prostate-Specific Antigens by Immunoaffinity-Mass Spectrometry. medRxiv, 2025.2006.2026.25330371 (2025).

28. Z. Fu, Y. Rais, D. Dara, D. Jackson, A. P. Drabovich, Rational Design and Development of SARS-CoV-2 Serological Diagnostics by Immunoprecipitation-Targeted Proteomics. Anal Chem 94, 12990–12999 (2022).

29. K. H. D. Crawford et al., Protocol and Reagents for Pseudotyping Lentiviral Particles with SARS-CoV-2 Spike Protein for Neutralization Assays. Viruses 12, (2020).

30. F. Krammer, P. Palese, Advances in the development of influenza virus vaccines. Nat Rev Drug Discov 14, 167–182 (2015).

31. G. Singh et al., Population immunity to clade 2.3.4.4b H5N1 is dominated by anti- neuraminidase antibodies. medRxiv, 2026.2002.2010.26346014 (2026).

32. T. K. Tsang et al., Association between antibody titers and protection against influenza virus infection within households. J Infect Dis 210, 684–692 (2014).

33. K. L. Winarski et al., Vaccine-elicited antibody that neutralizes H5N1 influenza and variants binds the receptor site and polymorphic sites. Proc Natl Acad Sci U S A 112, 9346–9351 (2015).

34. T. Zuo et al., Comprehensive analysis of antibody recognition in convalescent humans from highly pathogenic avian influenza H5N1 infection. Nat Commun 6, 8855 (2015).

35. Y. Zuo et al., Complementary recognition of the receptor-binding site of highly pathogenic H5N1 influenza viruses by two human neutralizing antibodies. J Biol Chem 293, 16503–16517 (2018).

36. N. L. Kallewaard et al., Structure and Function Analysis of an Antibody Recognizing All Influenza A Subtypes. Cell 166, 596–608 (2016).

37. T. Zuo et al., Comprehensive analysis of antibody recognition in convalescent humans from highly pathogenic avian influenza H5N1 infection. Nat Commun 6, 8855 (2015).

38. D. T. Bui et al., Mass Spectrometry-Based Shotgun Glycomics Using Labeled Glycan Libraries. Anal Chem 94, 4997–5005 (2022).

39. D. T. Bui, E. N. Kitova, L. K. Mahal, J. S. Klassen, Mass spectrometry-based shotgun glycomics for discovery of natural ligands of glycan-binding proteins. Curr Opin Struct Biol 77, 102448 (2022).

40. H. Park et al., Mass Spectrometry-Based Shotgun Glycomics for Discovery of Natural Ligands of Glycan-Binding Proteins. Anal Chem 92, 14012–14020 (2020).

41. D. T. Bui et al., Absolute Affinities from Quantitative Shotgun Glycomics Using Concentration-Independent (COIN) Native Mass Spectrometry. ACS Cent Sci 9, 1374–1387 (2023).

42. L. Glaser et al., A single amino acid substitution in 1918 influenza virus hemagglutinin changes receptor binding specificity. J Virol 79, 11533–11536 (2005).

43. N. C. Wu et al., A complex epistatic network limits the mutational reversibility in the influenza hemagglutinin receptor-binding site. Nat Commun 9, 1264 (2018).

44. M. Rios Carrasco et al., Acquisition of specific human respiratory tract binding of 2.3.4.4b H5N1 hemagglutinins requires multiple mutations. bioRxiv, 2026.2004.2016.718875 (2026).

45. M. Rios Carrasco et al., The Q226L mutation can convert a highly pathogenic H5 2.3.4.4e virus to bind human-type receptors. Proc Natl Acad Sci U S A 122, e2419800122 (2025).

46. B. Dadonaite et al., Deep mutational scanning of H5 hemagglutinin to inform influenza virus surveillance. Plos Biol 22, e3002916 (2024).

47. Alberta Government. (Alberta Government, Alberta Government, 2025), vol. 2026.

48. Y. Shu, J. McCauley, GISAID: Global initiative on sharing all influenza data – from vision to reality. Eurosurveillance 22, 30494 (2017).

49. S. Chakraborty, K. Trihemasava, G. Xu, Modifying Baculovirus Expression Vectors to Produce Secreted Plant Proteins in Insect Cells. J Vis Exp, (2018).

50. S. M. Ghafoori et al., Structural characterisation of hemagglutinin from seven Influenza A H1N1 strains reveal diversity in the C05 antibody recognition site. Sci Rep 13, 6940 (2023).

51. C. Q. Huang et al., Computationally designed haemagglutinin with nanocage plug-and- display elicits pan-H5 influenza vaccine responses. Emerg Microbes Infect 14, 2511132 (2025).

52. S. Meier, S. Guthe, T. Kiefhaber, S. Grzesiek, Foldon, the natural trimerization domain of T4 fibritin, dissociates into a monomeric A-state form containing a stable beta-hairpin: atomic details of trimer dissociation and local beta-hairpin stability from residual dipolar couplings. J Mol Biol 344, 1051–1069 (2004).

53. A. H. Keeble et al., Approaching infinite affinity through engineering of peptide-protein interaction. Proc Natl Acad Sci U S A 116, 26523–26533 (2019).

54. F. J. Milder et al., Universal stabilization of the influenza hemagglutinin by structure- based redesign of the pH switch regions. Proc Natl Acad Sci U S A 119, (2022).

55. I. Berger, D. J. Fitzgerald, T. J. Richmond, Baculovirus expression system for heterologous multiprotein complexes. Nature biotechnology 22, 1583–1587 (2004).

56. C. Bieniossek, T. J. Richmond, I. Berger, MultiBac: multigene baculovirus-based eukaryotic protein complex production. Current protocols in protein science / editorial board, John E. Coligan … [et al.] Chapter 5, Unit 5 20 (2008).

57. Y. Rais, A. P. Drabovich, Identification and Quantification of Human Relaxin Proteins by Immunoaffinity-Mass Spectrometry. J Proteome Res 23, 2013–2027 (2024).

58. Z. Fu et al., Mapping Isoform Abundance and Interactome of the Endogenous TMPRSS2-ERG Fusion Protein by Orthogonal Immunoprecipitation-Mass Spectrometry Assays. Mol Cell Proteomics 20, 100075 (2021).

59. J. Zhang et al., Germ cell-specific proteins AKAP4 and ASPX facilitate identification of rare spermatozoa in non-obstructive azoospermia. Mol Cell Proteomics 22, 100556 (2023).

60. C. Schiza, D. Korbakis, K. Jarvi, E. P. Diamandis, A. P. Drabovich, Identification of TEX101-associated Proteins Through Proteomic Measurement of Human Spermatozoa Homozygous for the Missense Variant rs35033974. Mol Cell Proteomics 18, 338–351 (2019).

61. A. P. Drabovich et al., Multi-omics biomarker pipeline reveals elevated levels of Protein- glutamine Gamma-glutamyltransferase 4 in seminal plasma of prostate cancer patients. Mol Cell Proteomics 18, 1807–1823 (2019).

62. T. D. Karakosta, A. Soosaipillai, E. P. Diamandis, I. Batruch, A. P. Drabovich, Quantification of Human Kallikrein-Related Peptidases in Biological Fluids by Multiplatform Targeted Mass Spectrometry Assays. Mol Cell Proteomics 15, 2863–2876 (2016).

63. Z. Fu, Y. Rais, D. Dara, D. Jackson, A. P. Drabovich, Rational Design and Development of SARS-CoV-2 Serological Diagnostics by Immunoprecipitation-Targeted Proteomics. Analytical Chemistry 94, 12990–12999 (2022).

64. P. I. Kitov, L. Han, E. N. Kitova, J. S. Klassen, Sliding Window Adduct Removal Method (SWARM) for Enhanced Electrospray Ionization Mass Spectrometry Binding Data. J Am Soc Mass Spectrom 30, 1446–1454 (2019).

65. B. Zakeri et al., Peptide tag forming a rapid covalent bond to a protein, through engineering a bacterial adhesin. Proc Natl Acad Sci U S A 109, E690–697 (2012).

66. A. Punjani, J. L. Rubinstein, D. J. Fleet, M. A. Brubaker, cryoSPARC: algorithms for rapid unsupervised cryo-EM structure determination. Nat Methods 14, 290–296 (2017).

67. T. Bepler et al., Positive-unlabeled convolutional neural networks for particle picking in cryo-electron micrographs. Nat Methods 16, 1153–1160 (2019).

68. E. C. Meng et al., UCSF ChimeraX: Tools for structure building and analysis. Protein Sci 32, e4792 (2023).

69. T. I. Croll, ISOLDE: a physically realistic environment for model building into low- resolution electron-density maps. Acta Crystallogr D Struct Biol 74, 519–530 (2018).

70. P. Emsley, B. Lohkamp, W. G. Scott, K. Cowtan, Features and development of Coot. Acta Crystallogr D Biol Crystallogr 66, 486–501 (2010).

71. D. Liebschner et al., Macromolecular structure determination using X-rays, neutrons and electrons: recent developments in Phenix. Acta Crystallogr D Struct Biol 75, 861–877 (2019).

72. V. B. Chen et al., MolProbity: all-atom structure validation for macromolecular crystallography. Acta Crystallogr D Biol Crystallogr 66, 12–21 (2010).

73. U.S. Food and Drug Administration. (U.S. Food and Drug Administration, U.S. Food and Drug Administration, 2024).

74. Health Canada. (Health Canada, Health Canada, 2025).

75. W. Li, L. Jaroszewski, A. Godzik, Tolerating some redundancy significantly speeds up clustering of large protein databases. Bioinformatics 18, 77–82 (2002).

76. K. Katoh, J. Rozewicki, K. D. Yamada, MAFFT online service: multiple sequence alignment, interactive sequence choice and visualization. Brief Bioinform 20, 1160–1166 (2019).

77. S. Kuraku, C. M. Zmasek, O. Nishimura, K. Katoh, aLeaves facilitates on-demand exploration of metazoan gene family trees on MAFFT sequence alignment server with enhanced interactivity. Nucleic Acids Res 41, W22–28 (2013).

78. S. Capella-Gutiérrez, J. M. Silla-Martínez, T. Gabaldón, trimAl: a tool for automated alignment trimming in large-scale phylogenetic analyses. Bioinformatics 25, 1972–1973 (2009).

79. S. Guindon et al., New algorithms and methods to estimate maximum-likelihood phylogenies: assessing the performance of PhyML 3.0. Syst Biol 59, 307–321 (2010).

80. F. D. Ciccarelli et al., Toward automatic reconstruction of a highly resolved tree of life. Science 311, 1283–1287 (2006).

